# Extreme male reproductive skew limits effective population size in American Standardbred horses

**DOI:** 10.64898/2026.01.03.697495

**Authors:** Robin S. Waples, Anas M. Khanshour, E. G. Cothran

## Abstract

Standardbred horses are widely used for harness racing in North America, Europe, Australia, and New Zealand. In the U.S., two gaits (Pacers and Trotters) are genetically differentiated at a level typical for different horse breeds. We used the complete pedigree for all registered American Standardbred foals born 1970-2014 (>200K Trotters and >500K Pacers) to calculate annual and lifetime reproductive skew and its consequences for the effective number of breeders per year (*N_b_*) and effective size per generation (*N_e_*). With rare exceptions, females can only produce 0 or 1 foal per year, which limits annual and lifetime variance in offspring number. These constraints do not apply to males: with humans (rather than horses) deciding who breeds with whom, and how often, the most prolific stud regularly sires well over 100 foals each year and often over 1000 across its lifetime. Male reproductive skew increased over time in both gaits, with the most prolific studs producing 20% or more of all foals sired by members of their birth cohort. Although each gait in the U.S. is represented by tens of thousands of adult horses at any given time, male *N_e_* is typically in the low hundreds, which drives the overall *N_e_*/*N* ratio to or below 0.01—a value often considered to be “tiny.” *N_e_*/*N* ratios this low have rarely been reported for large mammals and generally are thought to apply primarily to some marine species with very high fecundity. When *N_e_*/*N* is this low, even large populations can experience substantial erosion of genetic diversity.

## Introduction

The Convention on Biological Diversity (CBD), which was initiated at the 1992 Earth Summit in Rio de Janiero, has been ratified by all United Nations members except the United States. Curiously, given that genetic diversity ultimately is the foundation of all biodiversity, until recently the CBD said little specifically about conservation of genetic diversity in the wild (Laikre et al. 2010a). That changed with the CBD’s post-2020 global biodiversity framework, in which the Parties committed to monitoring genetic diversity for all species (CBD 2022; Carroll et al. 2023). Furthermore, the 2022 Kunming-Montreal Global Biodiversity Framework identified effective population size (*N_e_*) as a key metric for tracking within-population genetic diversity (Hoban et al. 2022; Mastratta-Yanes et al. 2024). *N_e_* determines the rate of random genetic drift and its consequences (rates of allele frequency change, loss of genetic diversity, and increase in inbreeding).

The rate of genetic drift is inversely, and non-linearly, related to *N_e_*. All else being equal, therefore, larger populations have larger *N_e_* and lower rates of genetic drift. In the real world, however, things are seldom that simple. Adult census size (*N_A_*) defines the number of potential parents and generally sets an upper limit to how large *N_e_* can be, but given that constraint the factor that has the largest influence on the *N_e_*/*N_A_* ratio is variation among individuals in number of offspring they produce. A handy point of reference is what is known as a Wright-Fisher “ideal” population, in which each individual has an equal opportunity to pass genes on to subsequent generations. In such a population, random mating and random reproductive success produce an approximately Poisson distribution of offspring number, in which the variance in offspring number approximately equals the mean and *N_e_*≈*N_A_*. In almost all real populations, however, the variance exceeds the mean (hence is overdispersed), with the result that *N_e_*<*N_A_*.

How much smaller than *N_A_* can effective size be? Despite active debate spanning several decades, no consensus has emerged in the scientific literature. Nunney (1993, 1995; Nunney and Elam 1994) has argued on theoretical grounds that *N_e_*/*N_A_* should generally be around 0.5 and that special considerations are required to produce a ratio as small as 0.1. At the other extreme, others (e.g., Hedgecock and Pudovkin 2011; Hauser and Carvalho 2008) have argued for a model of “sweepstakes” reproductive success, whereby most families produce no surviving offspring, while a few hit the jackpot and are genetically responsible for most of the next generation. The sweepstakes hypothesis requires high fecundity to allow a small number of individuals to each produce many offspring. Empirical support for this hypothesis can be found in several published genetic studies that estimate “tiny” *N_e_*/*N_A_* ratios (<0.01 or <0.001), primarily for marine species with high fecundity (Hauser and Carvalho 2008). But estimated *N_e_*/*N_A_* ratios are subject to a variety of potential biases (Waples 2016), and some other studies of high-fecundity species have produced estimates of *N_e_*/*N_A_* in the ‘normal’ range predicted by Nunney (Waples et al. 2018a; Jones et al. 2019).

Based on their life history traits, large mammals would seem to be unlikely candidates to exhibit sweepstakes reproductive success and a tiny *N_e_*/*N_A_* ratio. Females can typically only produce 0 or 1 (rarely 2) offspring per year, which severely constrains how large the variance in lifetime reproductive output (var(*LRO*)) can be. In some species (e.g., elephant seals (LeBoeuf and Reiter 1988), bighorn sheep (Coltman et al. 2002)), a few males control harems of many females. However, a) maximum family sizes are orders of magnitude smaller than for high-fecundity marine species, and b) few males can maintain large harems for many years—both of which limit how large var(*LRO*) can be.

But what if reproduction (who gets to mate, with whom, and how often) is controlled not by the animals themselves, but by humans? Each year, human intervention in reproduction occurs for countless species on a gigantic global scale (Laikre et al. 2010b). Artificial propagation programs have a variety of different and often overlapping goals, including food production, increasing harvest opportunities, conservation, and genetic improvement. These interventions can obviously affect the distribution of reproductive success, and a considerable literature exists regarding the potential and actual genetic consequences of artificial propagation (Ryman and Laikre 1991; Mitra 2001; Frankham 2008; Kitada 2018).

One thing these disparate efforts have in common is that decisions about reproduction are made in the context of overall program goals. Consider now a different model, where choices about reproduction are distributed broadly within the human community. For example, a recent report (Equo 2017) estimated that there are 2 million horse owners in the US alone, of which about 2% (40,000) own racehorses. Each year for each mare, a decision is made whether to breed the mare and, if so, to which stallion. Past performance by potential studs (in racing and breeding) and economic factors (including stud fees) are of course important considerations, but such decisions can also be driven by less tangible factors. This can lead to patterns similar to those that have been documented for choices of baby names: parents tend to favor names that have recently risen in popularity, in the same way that momentum traders in the stock market base decisions on recent price trends (Gureckis and Goldstone 2009). The degree to which individual female racehorses can dominate reproduction is still limited by biology, but the same constraints do not apply to males. In a system that operates in this fashion, how large can male skew be, and how low the *N_e_*/*N_A_* ratio?

Here we consider this question for American Standardbred horses, the breed used for harness racing. The Standardbred registry dates back only to 1871, but bloodlines for the breed originated in English Thoroughbreds. Standardbred horses use two different gaits: pacing and trotting. The gaits define subpopulations that are not entirely isolated but are characterized by moderate genetic differences (*F_ST_* ≈ 0.08; Esdaile et al. 2022) comparable to those typically found between different horse breeds. Our analyses focus on the complete pedigree for almost three-quarters of a million Trotter and Pacer foals born in the 45-year period from 1970 to 2014. We show that, within each gait: a) several indices of both male and female reproductive skew have increased over time; b) the most prolific studs sire over 100 foals each year and over 1000 over their lifetimes; and c) male *N_e_* is less than 1%, and overall *N_e_* less than 2%, of the number of adults.

## Materials and methods

### Pedigree data

Pedigree data supplied by U.S. Trotting Association (USTA) contained detailed information for every registered foal born 1970-2014, including birth year, gender, gait and parents. This dataset had 733,475 records, with 517,692 assigned as Pacers and 215,783 as Trotters (Table S1). Because about a third of all foals never race, we relied on the sire’s gait for the foals with no gaiting record. After removing foals with unknown or missing gender and those with parents >30 years of age (which occurred primarily in the earlier cohorts), we arrived at the following datasets used in all subsequent analyses: 215467 Trotter foals [108314 (50.3%) females and 107153 (49.7%) males] and 526118 Pacer foals [263125 (50.01%) females and 262993 (49.99%) males].

### Population demography

Most Standardbred foals are born in late winter or spring, so we analyzed the demographic data using the discrete-time, seasonal birth-pulse model (Caswell 2001), with age indexed by *x.* Individuals of age *x* produce an average of *b_x_* offspring and then survive to age *x*+1 with probability *v_x_*. Age at maturity (first age with *b_x_*>0) occurs at age α and maximum age is ω. Cumulative survival through age *x* is 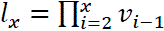, with *l*_1_=1. If population size is constant and if each birth cohort consists of *N_1_* foals, the expected number of individuals of age *x* alive at any given time is *E*(*N_x_*)=*N_1_l_x_*. Expected total census size is *E*(*N_T_*)=∑*E*(*N_x_*)=*N_1_Σ l_x_*, and expected total adult population size is 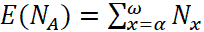.

Our pedigree data did not contain information about total population size—only those parents that actually produced offspring. To approximate total and age-specific population size, we used cumulative survivorship data (*l*_*x*_) for saddle horses shown in Figure 3 of Tapprest et al. (2017). For each gait, we set *N_1_* equal to the mean number of foals produced per year, and for the year (1969) that was immediately prior to our earliest pedigree data we estimated the stable age distribution as *N_x_* = *N_1_l_x_*. Each subsequent year, we used as *N_1_* the actual number of foals produced and propagated those values forward according to the survivorship schedule. Resulting estimates of *N_T_* and *N_A_* each year for both gaits are in Table S2. Raw estimates of age-specific fecundity were calculated from the empirical data, and these were rescaled to values expected in a constant population by using the stable-*N* relationship that ∑*b_x_l_x_*=2. Foals are registered at various times during their first year, so *b_x_* was scaled to production of offspring that survive to age 1.

Standard life tables include data for age-specific survival and fecundity, but we also are interested in a third age-specific vital rate, *V*_*k*,*x*_, which is the variance in offspring number (*k*) around the mean for individuals of age *x* (*b_x_*). We also consider the parameter *ϕ*_*x*_ = *V*_*k*,*x*_/*b*_*x*_, which is the ratio of variance to mean offspring number for individuals of age *x*. Crow and Morton (1955) called *φ* the “Index of Variability,” and its influence on effective size is apparent from Equation 1 below.

Assuming constant population size, expected generation length (*G* = average age of parents) can be calculated from vital rates as *E*(*G*) = ∑*xb_x_l_x_*/∑*b_x_l_x_*. From the actual pedigree, we estimated *G* each year in two ways. First, a static estimate *G_1_* was obtained by taking the average age of parents for all foals born each year. Second, a dynamic estimate *G_2_* was obtained by computing the average age at which all members of a cohort subsequently produced offspring. Thus, *G_1_* reflects ages of parents reproducing in a fixed time period, while *G_2_* is forward looking and reflects reproduction in the future by members of a cohort. In a population with constant *N* and stable age structure, *G_1_* and *G_2_* should be the same on average, but otherwise they can differ. Both *G_1_* and *G_2_* were computed separately for males and females, with the overall generation estimate being the mean across both sexes.

### Effective population size

In iteroparous species like horses, two types of effective size are important: *N_e_* = effective population size per generation and *N_b_* = the effective number of breeders in one year or breeding cycle. *N_e_* per generation is more important for long-term evolutionary processes, but *N_b_* provides more direct insight into the effects of the breeding system each year. Using the pedigree data described above, we calculated *N_b_* by focusing on the parents of foals produced each year, and we calculated *N_e_* by focusing on lifetime offspring produced by foals in each yearly birth cohort. We did this separately for Pacers and Trotters.

### *N_b_* per year

*N_b_* is determined by the mean (̄*k*) and variance (*V_k_*) among individuals in the number of offspring produced in a given year. Typically, *N_b_* is calculated using the standard formula for inbreeding effective size with separate sexes (Lande and Barrowclough 1987; Caballero 1994), which also requires one to know the number of potential parents, *N*:

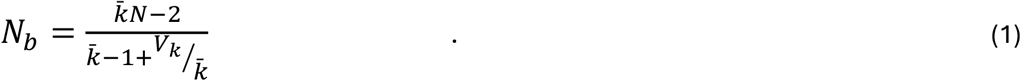

Inbreeding effective size, as opposed to variance effective size, is the most meaningful way to calculate *N_b_* because it provides information about the parents and is not affected by the total number of offspring produced, or by the size of the sample of offspring (Waples 2002). In this standard formulation, ̄*k* and *V_k_* are calculated across all mature individuals, including those that do not actually produce any offspring in the time period of interest. In the pedigree, however, we only have information about the parents of every foal that was actually born, so we lack information about any sires or dams that produced no foals in a given year. Thus, it is not possible to calculate overall *N* from the pedigree data alone. Fortunately, Waples and Waples (2011) showed that knowing the total number of parents is not essential; inbreeding effective size can be calculated based entirely on the vector of *k_i_* values, which is the numbers of offspring produced by each parent:

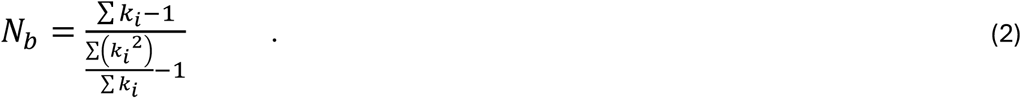

Parents that produce no offspring (*k_i_* = 0) make no contribution to any of the summation terms in Equation 2 and hence can be safely ignored. Equation 2 is used for each sex separately and combined to obtain an overall *N_b_* using Wright’s (1938) adjustment for sex ratio:

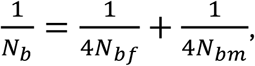

where *N_bf_* and *N_bm_* are the effective numbers of male and female breeders, respectively. Male *N_bm_* was calculated each year by allocating all the foals to sires to generate the vector of *k_i_* values. Because each mare that reproduces has a single foal each year (twins are very rare and ignored here), each foal has a different mother, in which case female *N_bf_* is infinitely large (Waples and Antao 2014). In that case, overall *N_b_* is given by:

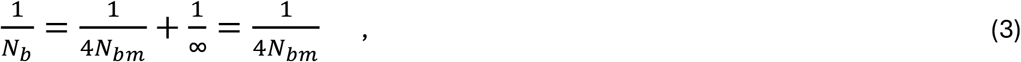

which means that overall *N_b_* for a reproductive cycle is 4 times *N_bm_*. We also identified males each year that produced unusually large numbers of offspring (>100), and we calculated the fraction of total yearly foal production attributed to the most prolific sires.

### *Ne* per generation

In iteroparous species with overlapping generations, effective size per generation is calculated as (Hill 1972, 1979):

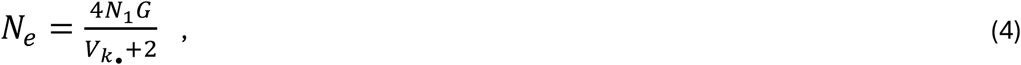

where *N_1_* is the number of offspring (foals) produced each year, and *V*_*k*•_ is the variance in the lifetime number of offspring produced by the *N_1_* individuals in a cohort. We calculated *V*_*k*•_, *G*, *N_1_*, and *N_e_* separately for males and females from the pedigree data and used Wright’s sex-ratio formula to calculate overall *N_e_*. We also evaluated both methods for estimating generation length (*G_1_*, *G_2_*).

Although lifetime ̄*k*_•_was nearly equal for males and females from each cohort, the number of foals produced per year changed over time, particularly in Pacers, and as a consequence ̄*k*_•_ varied across years. In contrast, Hill’s model assumes population size is constant, in which case lifetime ̄*k*_•_ for the population as a whole is 2. To account for variation in ̄*k*_•_, we scaled male and female *V*_*k*•_to values what would be expected when ̄*k*_•_= 2, using the following formula, adapted from (Waples 2002):

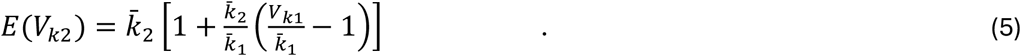

In Equation 5, ̄*k*_1_ is the observed mean lifetime number of offspring, ̄*k*_2_ is the target value (2) for a constant population, *V_k_*_1_ is the observed lifetime variance in reproductive success, and *V*_*k*2_ is the scaled variance in reproductive success, obtained by substituting the other values and solving the equation. Waples (2024) showed that under a model of random demographic stochasticity in *N* and ̄*k*_•_, this approach produced unbiased estimates of *N_e_*. As with the yearly *N_b_* data, for each cohort we calculated the fraction of all lifetime offspring produced by that cohort that was attributed to a single sire. This provides a useful index of over-dispersed variance in reproductive success, which has a large influence on *N_e_* per generation.

We used the *lm* function in R to test for significantly positive or negative temporal changes in key metrics. Evaluations of annual reproduction used time series with data for *n*=45 years, and evaluations of lifetime reproduction used time series with data for *n*=21 cohorts.

Variance in lifetime reproductive success (*V*_*k*•_) is enhanced (and *N_e_* correspondingly reduced) when persistent individual differences (Lee et al. 2011) exist—that is, when some individuals are consistently over time above or below average at producing offspring. Persistent differences can be quantified by computing correlations between an individual’s reproductive output at different ages. We did this in two ways. First, we calculated ρ_α,α+_ (Waples 2023), which is the Pearson correlation between offspring number at age at maturity (α) and for the rest of that individual’s lifetime. Because age at first reproduction is not fixed in Standardbreds, offspring number at age at maturity included all offspring produced for ages 3-5. Second, to provide more detailed insights into patterns of persistent differences, we calculated correlations of offspring production at ages separated by 1 to 15 years (the “age gap”). Both types of analysis were conducted separately for both sexes and both gaits, for all birth cohorts 1970-1990.

## Results

### Vital rates

Total numbers of registered Trotter foals per year in the period 1970-2014 ranged from 3197 to 6020 (mean = 4795), with peaks in the mid 1980s and mid 2000s and troughs around 1970, the mid 1990s, and in the most recent years (Table S1), but across all years there was no significant trend in foal production. Numbers of Pacer foals were higher (mean = 11504), starting around 10,000 in 1970, increasing to 18,000 by 1985 before monotonically declining to less than 5000 by 2014, leading to a highly significant overall decline (Table S5). The numbers of foals actually used in this study (Figure 1) were a few percent smaller. The sex ratio at birth was very close to 1:1 for both gaits, but approximately half of the male Standardbred foals are gelded each year [mean (range) for Trotters = 46.7% (37.0-55.0%); for Pacers = 55.7% (47.8-65.7%)], which restricts the pool of males available for breeding.

**Figure 1.**
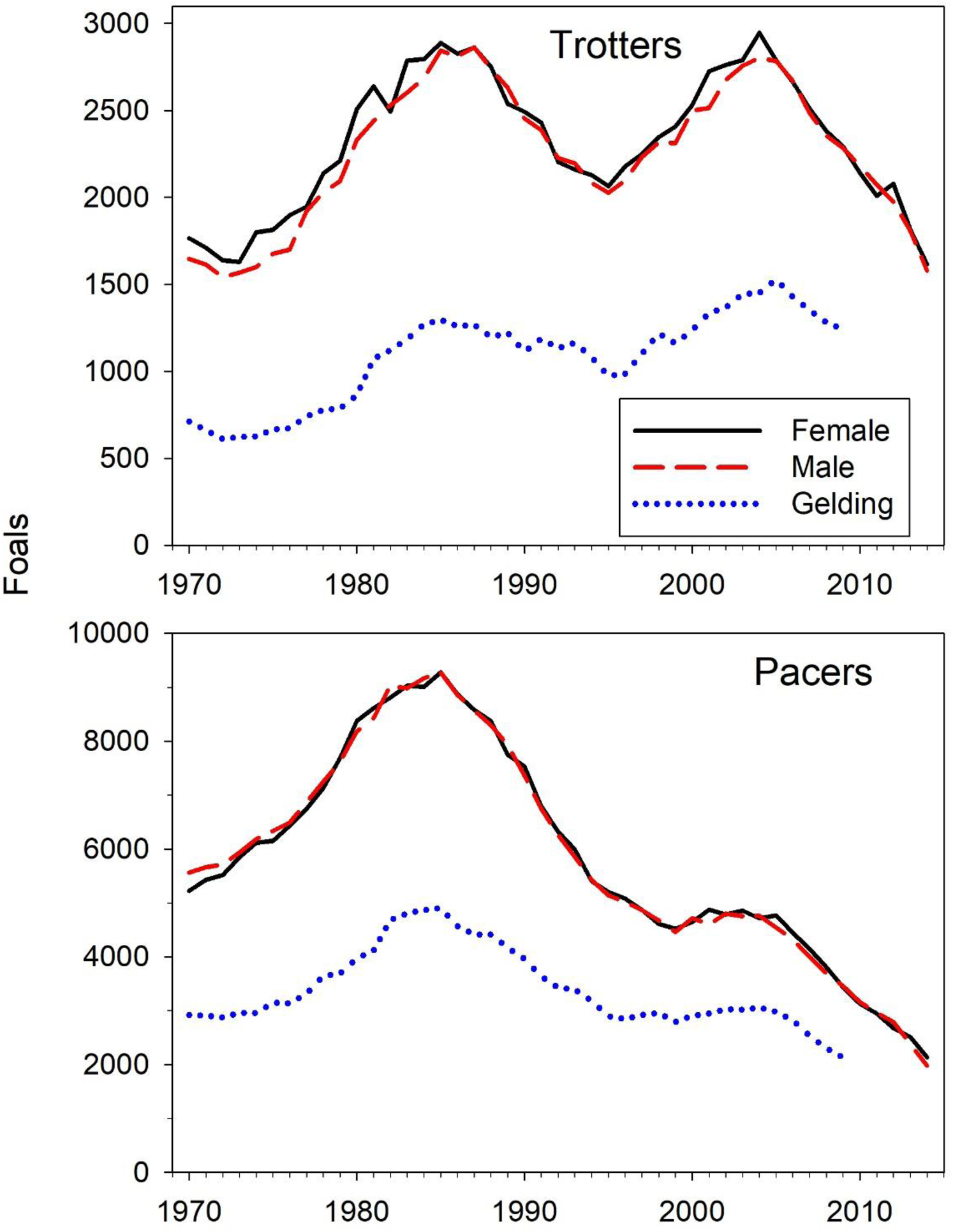
Number of registered foals per year by sex for Trotters (top) and Pacers (bottom. The number of males that were gelded is shown in dotted blue lines; data after 2009 are not complete.

Life tables (age-specific survival and fecundity) used for modeling Trotters and Pacers are given in Table S3. For both gaits, modeled survival was high (≥0.94) through age 12 and moderately high (>0.8) through age 24 before declining for the oldest ages. Maximum age was set at 30 as very few animals of either sex reproduced beyond their mid 20s. In both sexes in both gaits, age-specific fecundity (*b_x_*) peaked around age 10 before declining slowly during the teens and more rapidly after age 20 (Figure 2). Few mares gave birth after age 25, but a number of stallions were still active into their late 20s.

**Figure 2.**
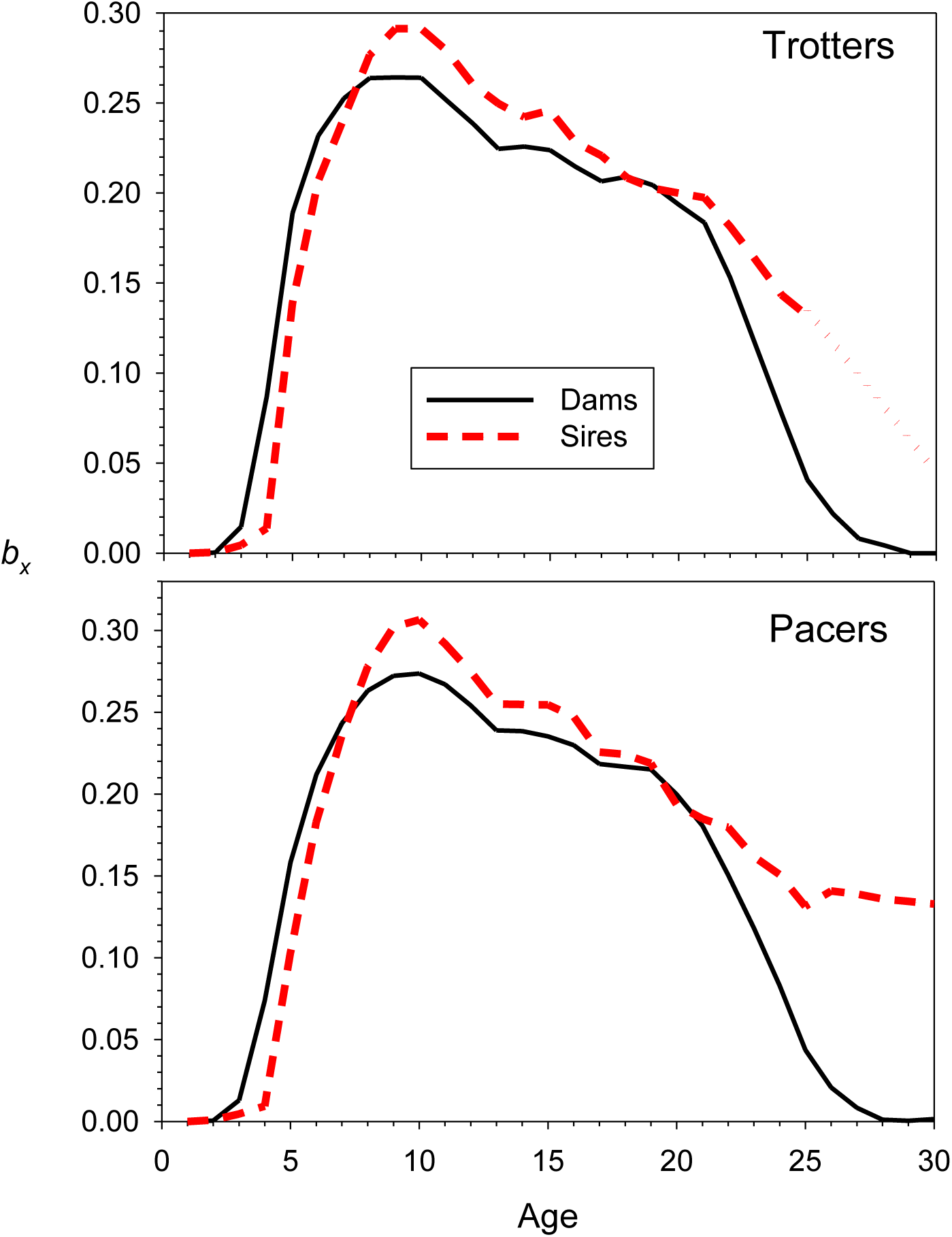
Age-specific fecundity (*b_x_*; from Table S3) for male and female Trotters (top) and Pacers (bottom). Values for Trotter males > age 25 (dotted red lines) are modeled based on sparse data.

**Figure 3.**
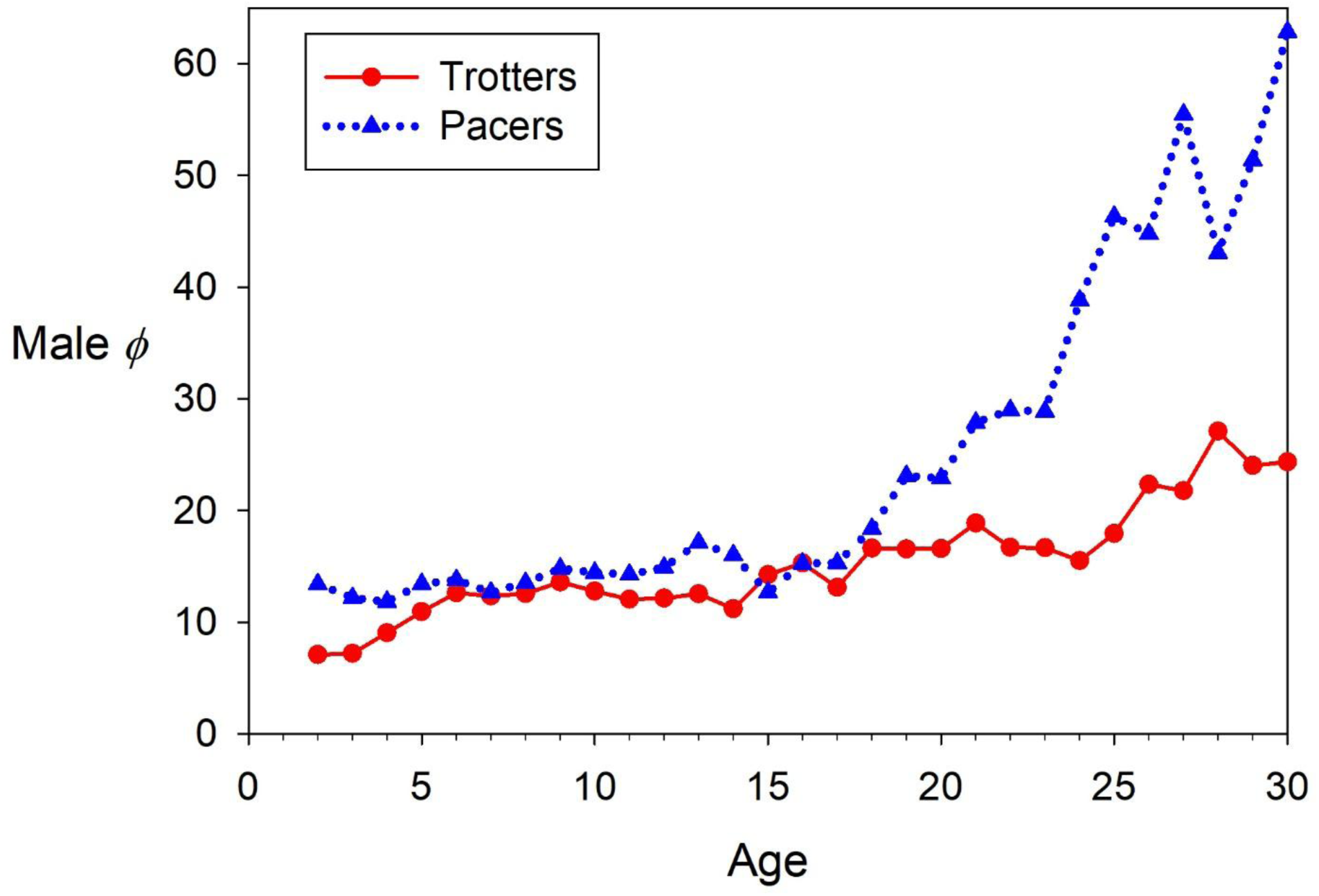
Age-specific vales of *φ* (ratio of *V*_*k*,*x*_ to *b_x_*) for male Trotters (solid red lines and red circles) and Pacers (dotted blue lines and blue triangles).

Patterns of the age-specific index of variability (*ϕ_x_*) differed dramatically between males and females. With very few exceptions (rare occurrence of twins), the only options for female Standardbred horses are to produce 0 or 1 offspring in a given year. In that case, *V*_*k*,*x*_ can never be as large as *b_x_*, so *ϕ* for females is always <1. In contrast, even for young male Trotters (<5 years of age), *ϕ_x_* was >5 and it increased steadily with age, averaging 12.4 for ages 6-14 and 16.2 for ages 15-25 (Figure 3). Male reproductive skew was even more extreme in Pacers: *ϕ_x_* was >10 for the youngest males, increased sharply after age 15, and exceeded 40 for ages ≥25—meaning that variance in annual offspring production was more than 40 times that expected under random reproductive success.

Across the full dataset, mean age at reproduction in Trotters was 11.2 for males and 10.5 for females; for Pacers the respective values were 11.4 and 10.8. Overall generation length is the mean across sexes, so *G* = 10.9 in Trotters and 11.1 in Pacers. As described in Methods, two methods were used to calculate generation length for specific time periods. Method *G*1 (mean age of parents reproducing each year) produced similar patterns in both gaits over the first 15 years of data: initial (1970) values were roughly 13 in males and 11 in females (so overall *G*1 was about 12 years initially), but these dropped continuously until reaching *G*1<10.5 in both sexes by 1985 (Figure 4). Subsequently, *G*1 in Pacers rose again and reached approximately the 1970 values for both sexes by the end of the time series. *G*1 for Trotter mares remained relatively stable after 1985, but the pattern was more variable for Trotter stallions.

**Figure 4.**
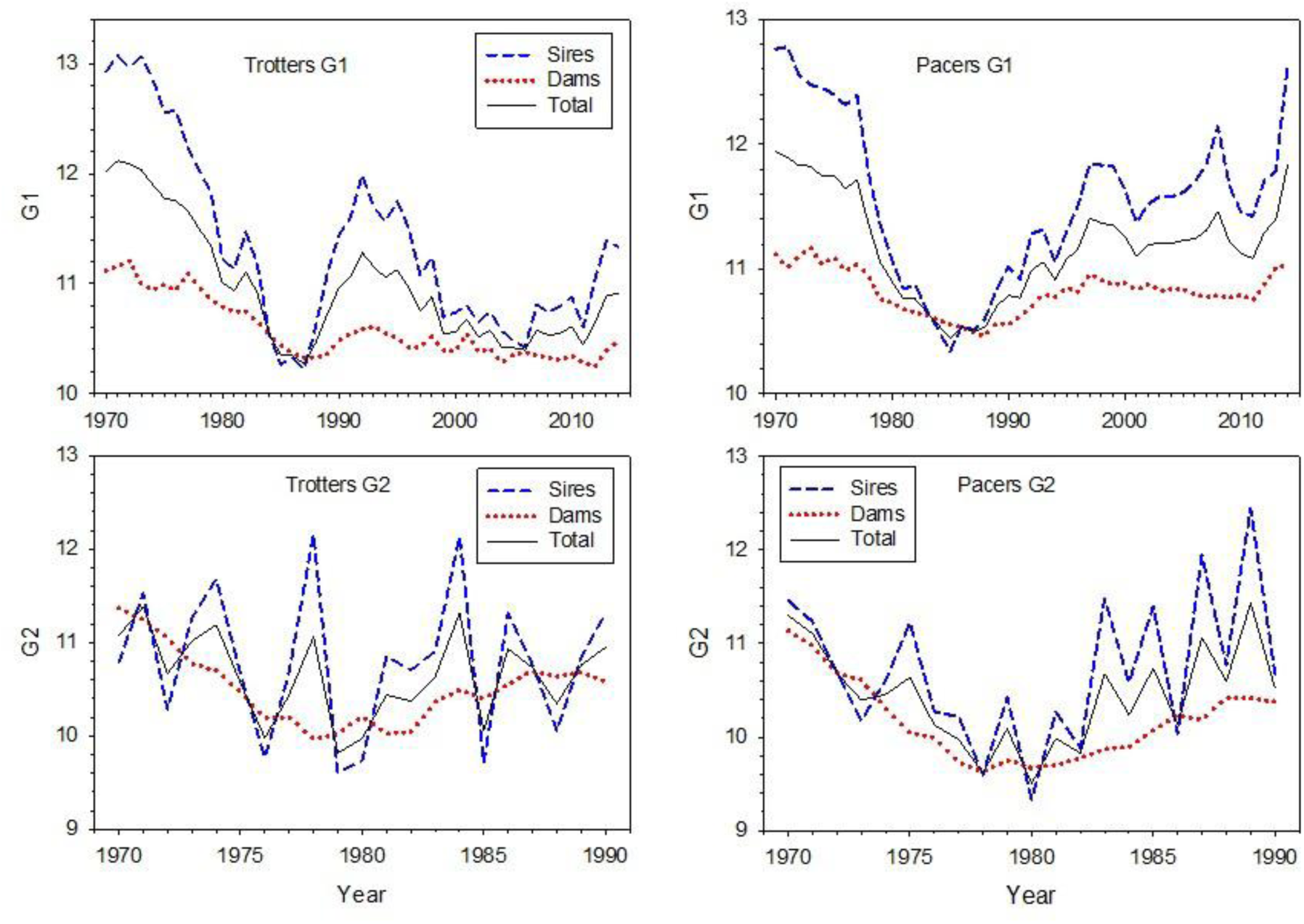
Generation length in Trotters (left panels) and Pacers (right panels) for males (dashed blue lines) and females (dotted red lines). Generation length is calculated using two methods. G1: mean age of parents of all foals born in each year. G2: mean age at which foals born in a yearly cohort subsequently produced offspring during their lifetime. Solid black lines (mean for males and females) are overall generation length.

Because *G*1 reflects reproduction by all adults alive within a given year (mostly >40,000 adults for Trotters and >80,000 for Pacers; Table S2), changes in generation length are relatively slow. In contrast, Method *G*2 tracks ages at future reproduction by members of a cohort, which number in the low thousands. Furthermore, as discussed below, individual males can dominate lifetime foal production within a cohort and hence have a disproportionate effect on *G*2. In both gaits, this results in a jagged, sawtooth pattern in male *G*2 (Figure 4). Of necessity, foal production is spread out among many more different females, leading to smooth and relatively small changes in female *G*2. Because of its relative stability, we used *G*_1_ for estimating *N*_e_ using Hill’s model.

### Annual reproduction

The fraction of adult females that foaled within a given year ranged from just under 10% to almost 28% and showed similar temporal patterns in both gaits, with cyclical changes but a net and highly significant decline (Figure S1; Table S5). In Trotters, twins occurred on average of 1.0 times per year, and this represented about 0.02% of the females. In Pacers, twins occurred on average of 3.4 times per year, and this represented about 0.03% of the females.

Across all ages, annual production of offspring by males was highly skewed and increasingly so over time in both gaits. The fraction of adult males producing at least one foal was never as high as 5% and dipped below 2% in the most recent years, showing a highly significant decline (Figure 5; Table S5). The most prolific Trotter stud never produced more than 88 foals in one year before 1978, but from 1980 to 2005 the biggest annual stud was called into service an average of 133 times (Figure 6; Table S4). In Pacers, maximum output by a single stud increased from 94 in 1970 to a peak of 235 in 1993. As a fraction of total foals, annual offspring production by the biggest stud significantly increased over time in both gaits, but more rapidly for Pacers than Trotters (Figure 7; Table S5). By 2010, in both gaits, the most prolific stud typically sired about 2.5-3% of all foals.

**Figure 5.**
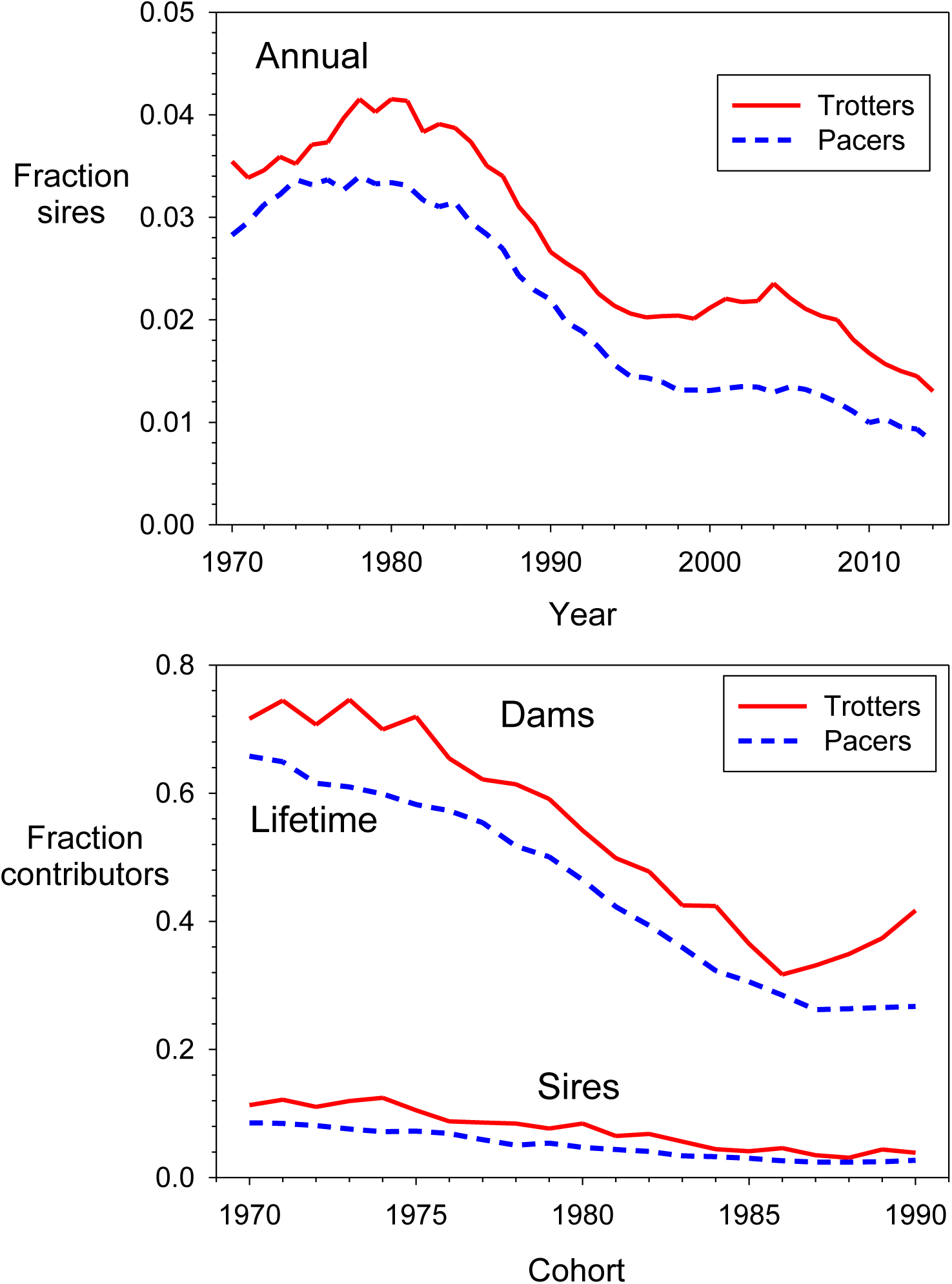
Fraction of all adults contributing to reproduction. Top: fraction of adult horses that produced at least one foal each year. Bottom: fraction of members of a newborn cohort that produced at least one foal during their lifetime. Solid red lines are data for Trotters and dashed blue lines are for Pacers.

**Figure 6.**
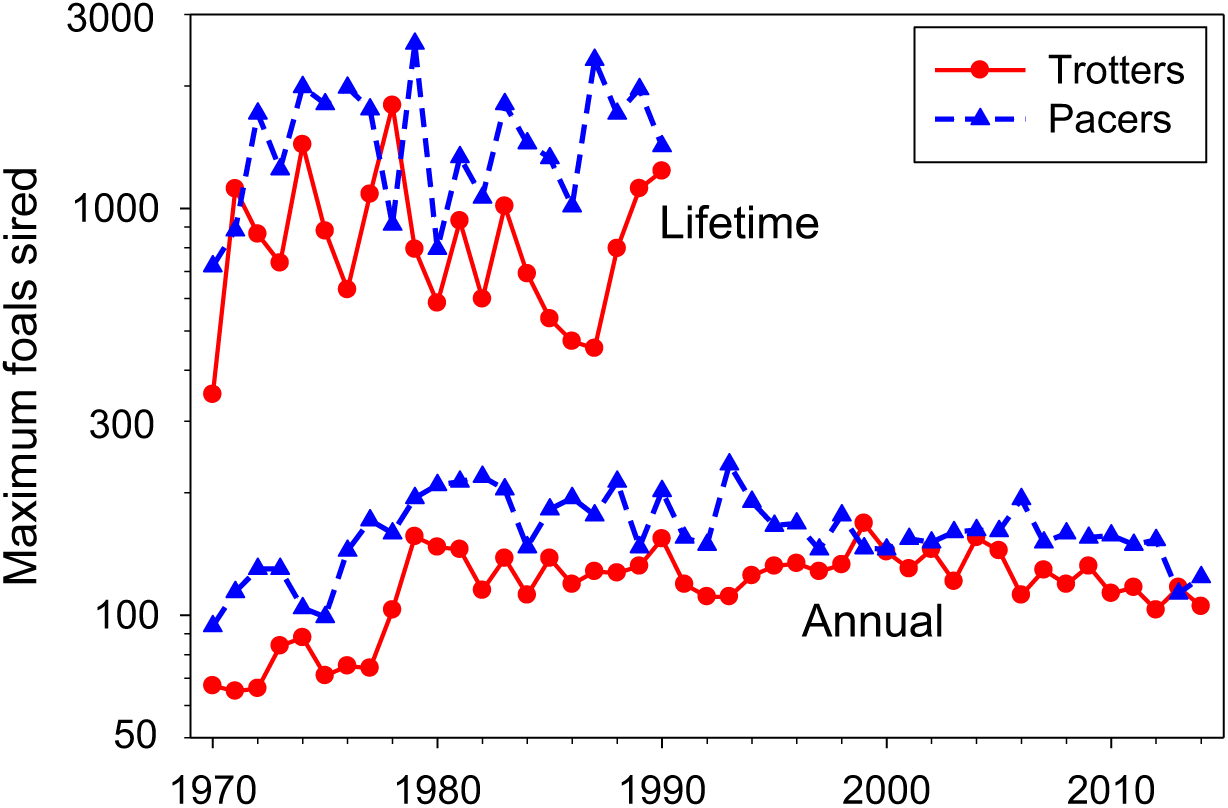
Maximum production of foals by a single male, annually and lifetime for cohorts born in the year indicated. Solid red lines are data for Trotters and dashed blue lines are for Pacers. Note log scale on the Y axis. Lifetime data are not complete for cohorts born after 1990.

**Figure 7.**
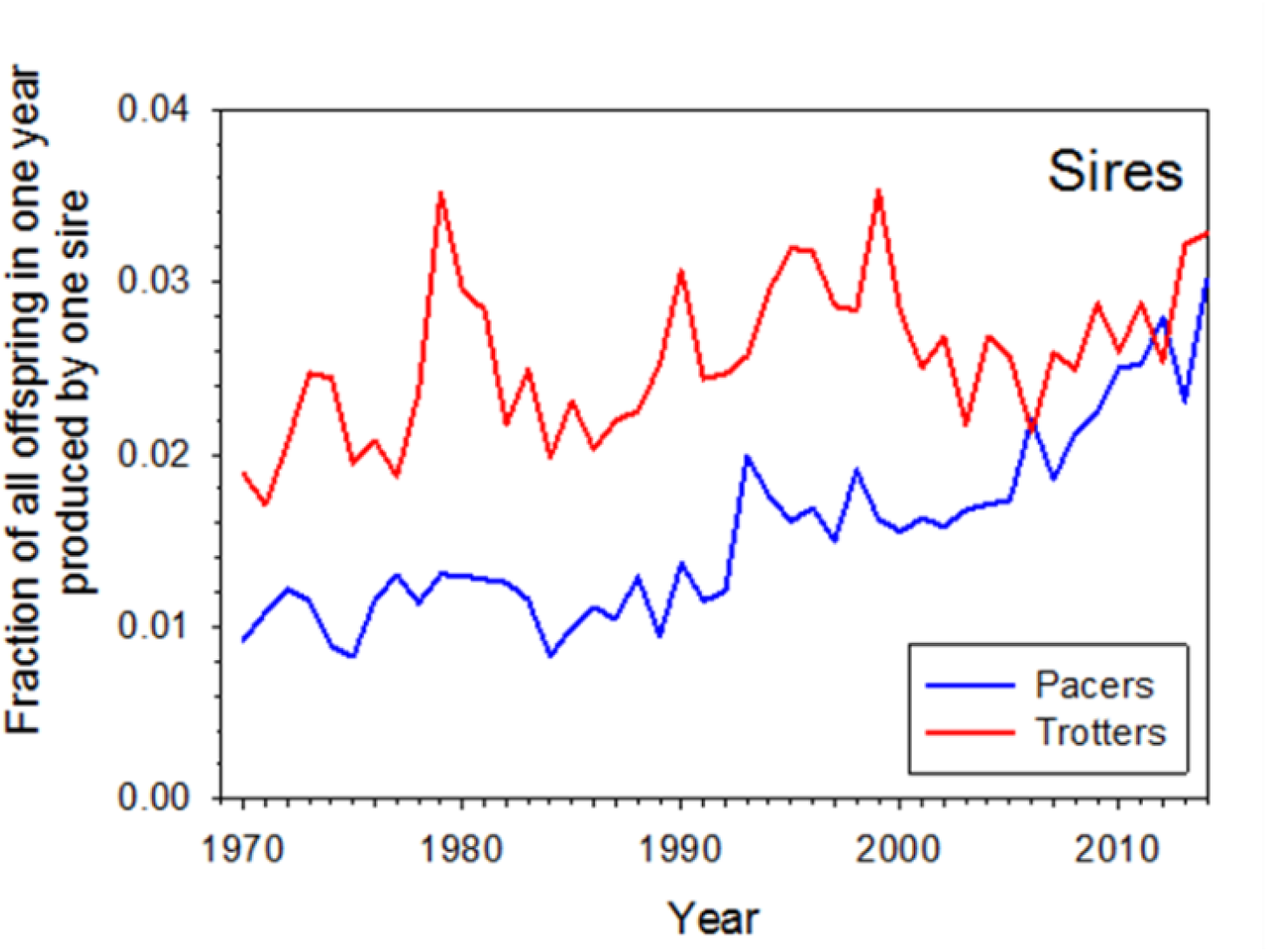
The relative influence of the most dominant male each year increased over time in both breeds, but more so in Pacers than Trotters.

The annual effective number of breeders (*N_b_*) significantly declined over time in both gaits, with the steeper decline in Pacers affected by a decline in population size from a peak in the mid 1980s (Figure 8; Table S5). After about 2005, *N_b_* per year averaged less than 1000 for both gaits. As a consequence of increasingly strong male reproductive skew, the estimated ratio *N_b_*/*N_A_* also declined sharply and significantly in both gaits, from about 5-8% before 1980 to about 1-2% in the most recent years (Figure 8; Table S5).

**Figure 8.**
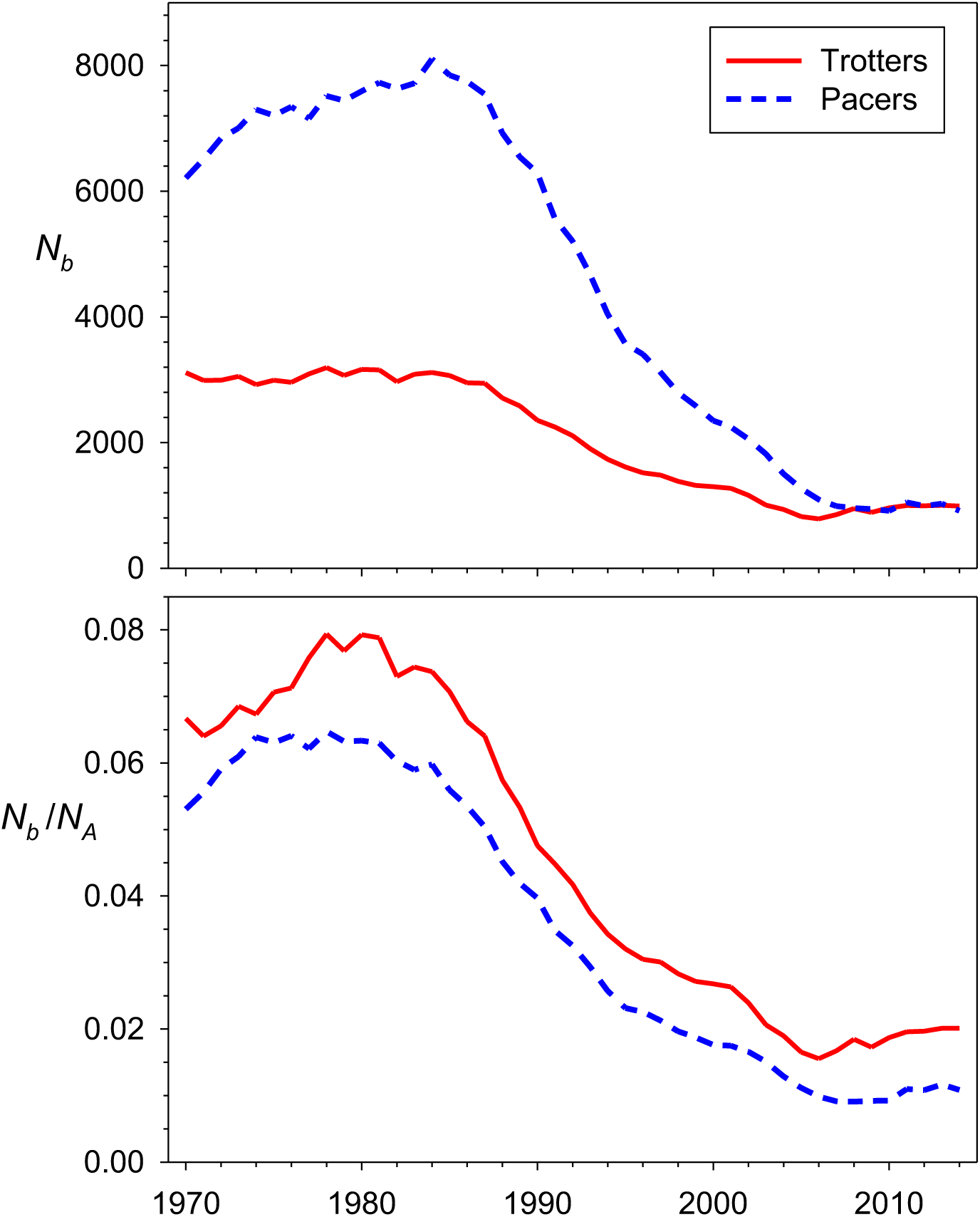
Pedigree-based *N_b_* (top) and *N_b_*/*N_A_* (bottom) in Trotters (solid red lines) and Pacers (dashed blue lines). In the lower panel, *N_A_* is estimated adult census size from Table S2.

**Figure 9.**
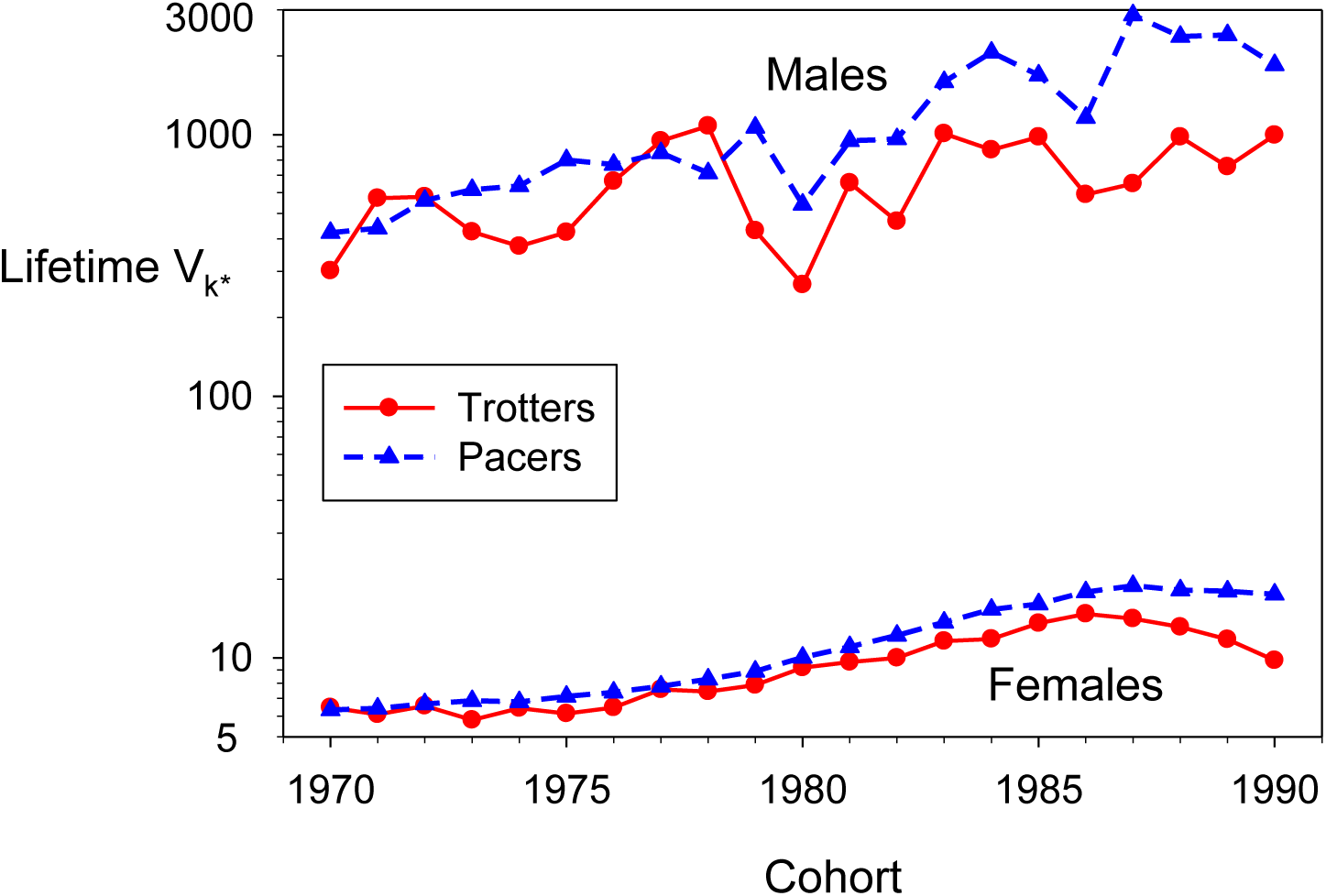
Lifetime variance in offspring number (*V*_*k*•_) for cohorts of males and females. *V*_*k*•_ is scaled to its expected value in a population of constant size. Solid red lines are data for Trotters and dashed blue lines are for Pacers. Note the log scale on the Y axis.

Across 45 years (1970-2014), foal production significantly declined in Pacers but showed no consistent trend in Trotters (Figure 1; Tables S1 and S5).

### Lifetime reproduction

Analyses of lifetime reproduction focus on cohorts of foals born 1970-1990, as some individuals in later cohorts were still reproductively active at the end of the data series.

Although annual reproduction of Standardbred mares is heavily constrained, a reproductive lifespan of over 20 years provides substantial opportunity for variation in lifetime offspring number. Within a cohort, maximum lifetime offspring production by mares ranged from 15 to 21 in Trotters and from 16 to 21 in Pacers, and oldest reproduction within cohorts occurred at ages 24-27 (Table S4). Of all the female foals born into a cohort, the fraction that produced at least one offspring in their lifetime significantly declined over time in both gaits (from about 70-75% in the early 1970s to 30-40% (Trotters) and 25-30% (Pacers) in cohorts born after 1985 (Figure 5; Table S5). These factors caused variance in female lifetime reproductive success to be overdispersed, with *V*_*k*•_ significantly increasing from about 5-6 in early cohorts (both gaits) to over 10 (Trotters) and over 15 (Pacers) in later cohorts (Tables S4 and S5). These values are many times the mean lifetime offspring number in a stable population.

Within a cohort of males, the fraction that sired at least one foal in their lifetime declined sharply and significantly over time in both gaits. In the earliest cohorts (1970-1974), an average of 11.8% of Trotter foals were lifetime contributors, while in the latest cohorts (1986-1990) the contributors had dropped to a mean of 3.9%; for Pacers, the respective numbers were 7.9% and 2.5% (Figure 5). Maximum lifetime foal production by Standardbred sires can be staggeringly high. In Trotters, the maximum stud within each cohort (that is, the one who won the Genghis Khan award) sired an average of over 800 foals (maximum = 1791); for Pacers, the mean was over 1500 foals, with a maximum of 2541 (Figure 6). For Trotters, these maximum studs were responsible for about 10% to 33% of all foals sired by their entire cohort (Figure 10). Although within each cohort the maximum Pacer stud typically produces more lifetime foals than the maximum Trotter stud (Figure 6), the fractional contributions of the Pacer studs were generally smaller (Figure 5) because Pacers considerably outnumber Trotters (Figure 1; Table S2). However, with the declining population size of Pacers, the proportional contribution of the maximum Pacer studs increased over time, reaching an average of 18.2% for the most recent 4 cohorts.

**Figure 10.**
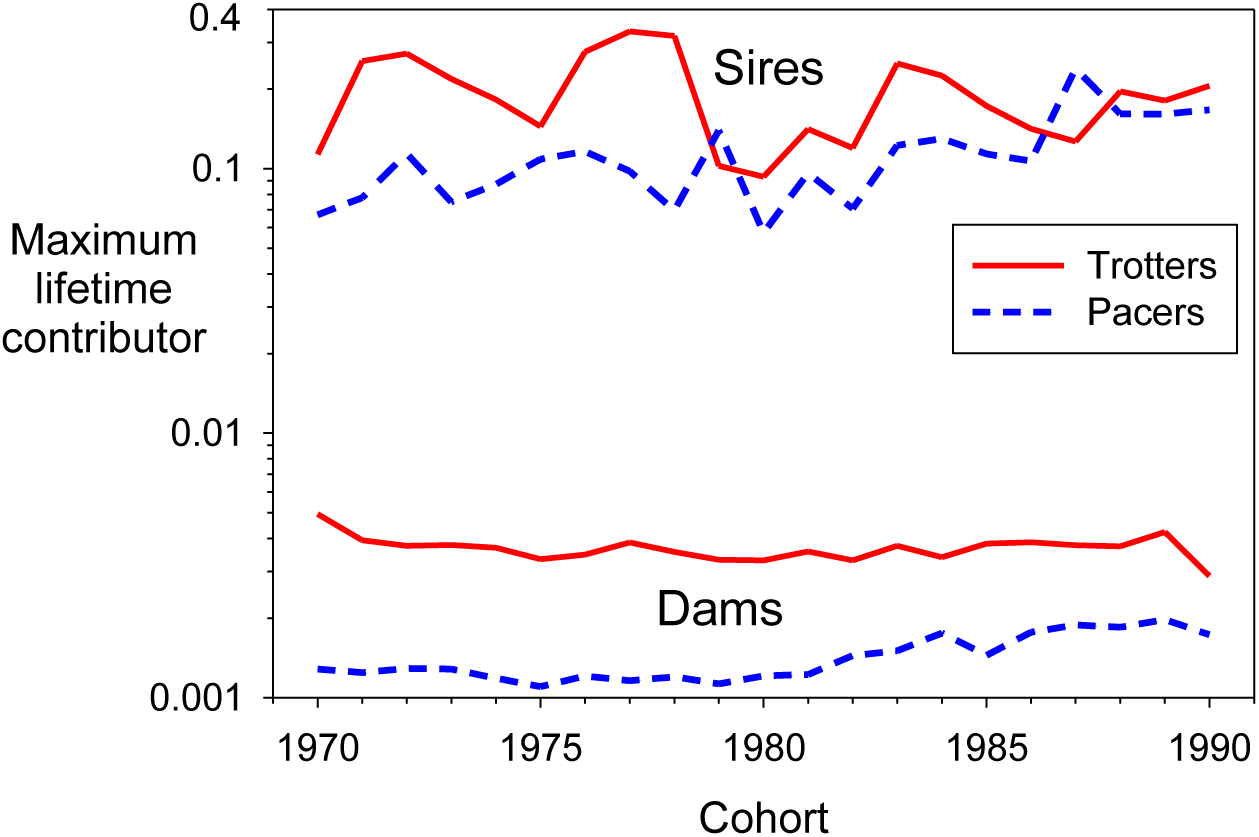
Maximum proportional lifetime contribution by individual sires and dams in each cohort. Values on the Y axis are the fraction of all offspring produced by members of a cohort that are attributed to a single individual. Solid red lines are data for Trotters and dashed blue lines are for Pacers. Note the log scale on the Y axis.

Some insights into factors that shape lifetime reproductive success of males can be gained by examining longitudinal data for individual sires. In Figure 11, foal production by age is tracked for two males within each gait: the male with the highest lifetime foal production (Sire A for Trotters and Sire C for Pacers), and the male who was responsible for the largest fraction of all foal production by his cohort (Sire B for Trotters and Sire D for Pacers). It is clear that lifetime foal production numbers this high require an individual male to consistently sire a large number of foals over a considerable period of time. Foal production by Sire A was uneven through age 10, but after that he sired a minimum of 77 foals in each of the next 16 years—still producing 79 at age 26. Sire C started earlier, having his first 100+ foal year at age 5 and maintained that level through age 20 before dropping off in his early 20s.

**Figure 11.**
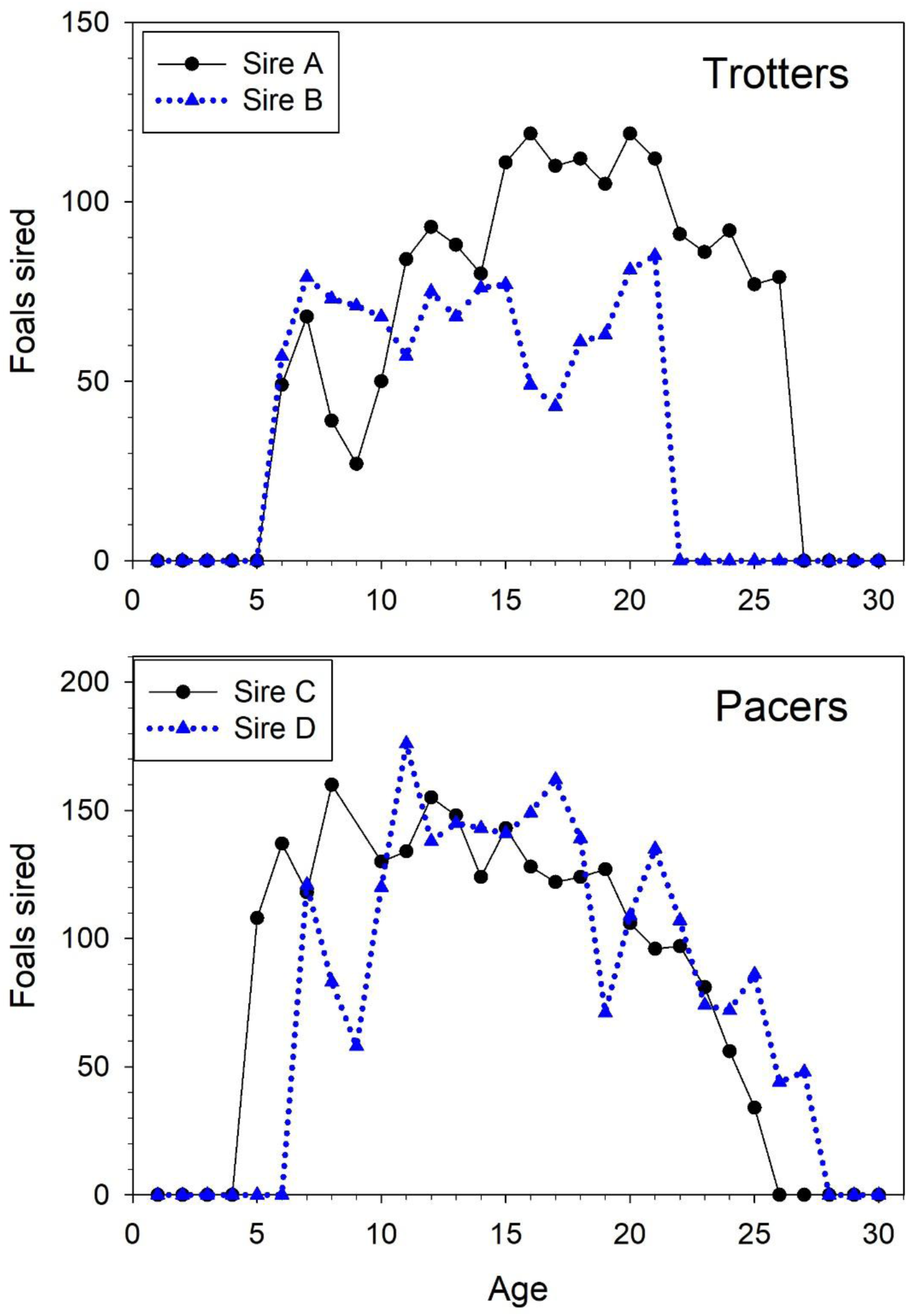
Number of foals sired by age for the most prolific male Trotters (top) and Pacers (bottom). Black lines and circles are results for the males in each breed that sired the most foals across its lifetime (1791 foals for sire A and 2541 for sire C). Blue lines and triangles are results for males that sired the largest fraction of foals produced by their entire cohort (33.1% for sire B and 23.9% for sire D). Mean age at reproduction was 17.0, 13.5, 13.7, and 16.0 for sires A-D, respectively, considerably higher than the overall means for male Trotters (11.2) and Pacers (11.4).

In the full datasets for both gaits, we also found strong evidence for persistent individual differences. Across 21 birth cohorts for males, median ρ_α,α+_ was 0.54 for Trotters and 0.51 for Pacers. Individual correlations were highest for consecutive years of reproduction (0.7-0.9) and were still high (0.4-0.6) for age gaps of 10 years (Figure S2). Somewhat surprisingly, we also found positive (albeit weaker) temporal correlations in reproductive success for females. For dams, median ρ_α,α+_ was 0.056 for Trotters and 0.057 for Pacers. Correlations for consecutive years of reproduction were >0.35 for females of both gaits; these correlations were still positive (∼0.1-0.2) for age gaps of 10 years (Figure S2).

The combination of a) consistently very high foal production by the maximum studs, and b) increasing incidence of null parents (those that never produce any offspring) resulted in a pattern of significantly increasing lifetime variance in offspring number (*V*_*k*•_) for males of both gaits (Figure 9; Table S5). These values are standardized to remove effects of mean offspring number and therefore represent a meaningful index of reproductive skew. For the most recent 5 cohorts, lifetime *V*_*k*•_ averaged almost 800 in Trotter males and over 2000 in Pacer males—compared to an expected value of 2 if foal production were randomly distributed among all males.

Female effective population size per generation (*N_e_*) declined about 15% in Trotters (from a harmonic mean of 9157 in early cohorts to 7790 in later ones) and by over 40% in Pacers (from 28046 to 16640) (Figure 12; declines were highly significant in both gaits, Table S5). Male *N_e_* in Pacers showed a steep (>2x) and highly significant decline over time (harmonic mean = 468 in first 5 cohorts vs 173 in latest 5 cohorts), whereas changes between early and late cohorts in Trotters (155 vs 145) were small and not significant. Overall *N_e_* (twice the harmonic mean for males and females) is driven primarily by the smaller male values, and temporal patterns of change were similar to those for males.

**Figure 12.**
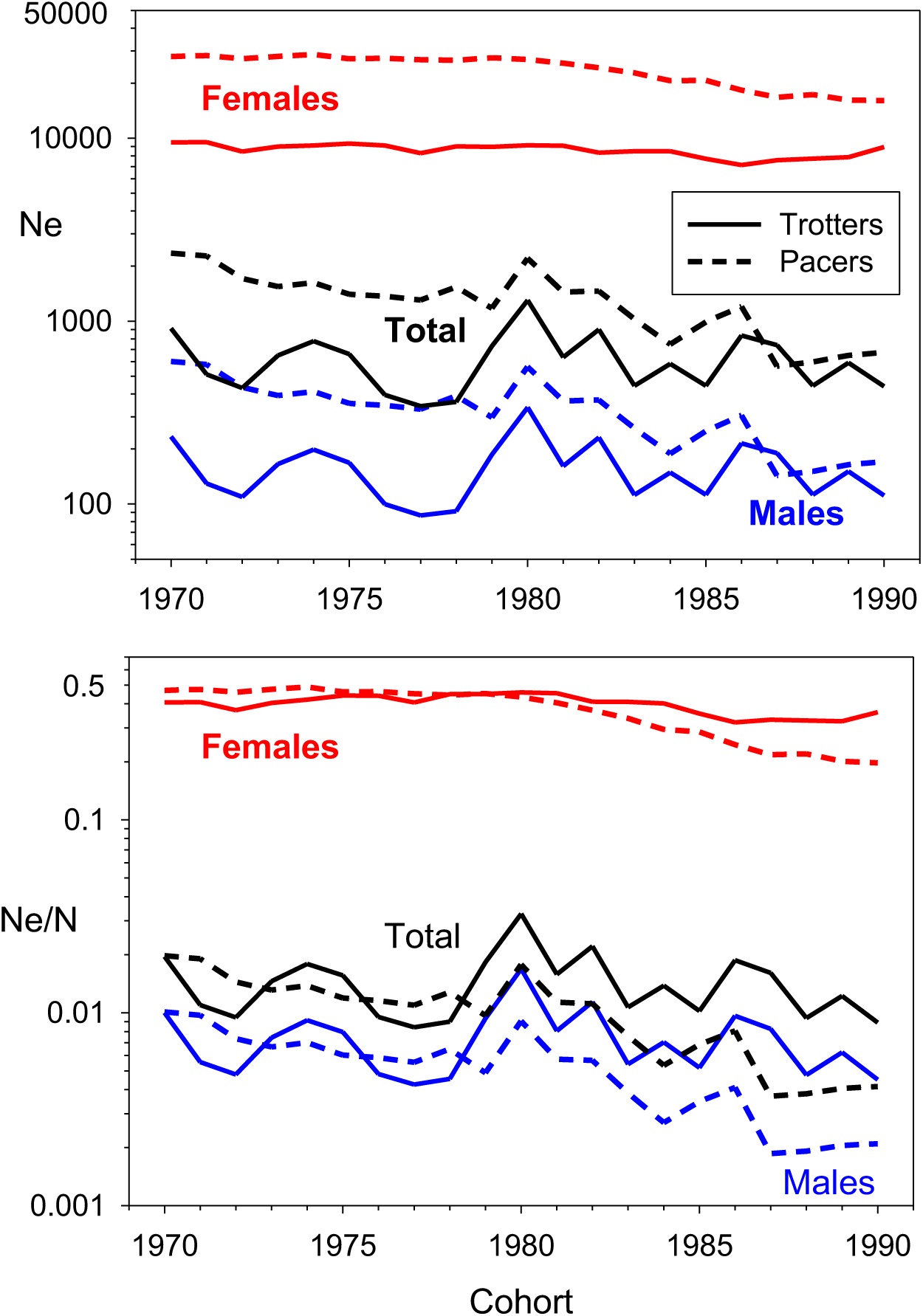
Pedigree-based effective population size per generation (*N_e_*) and the estimated *N_e_*/*N* ratio for each cohort of females (red), males (blue), and overall (black). Solid lines are data for Trotters; dashed lines are data for Pacers. In the lower panel, *N_A_* is estimated adult census size for from Table S2 for the year the cohort was born.

Context for understanding disparities in *LRO* for Standardbred horses can be provided by reference to the Gini index (Gini 1936), developed to quantify the degree of income inequality. The Gini index ranges from 0 (everyone gets the same amount of money) to 1 (Elon gets all the money). Most-recent Gini values by country range from 0.24 (Slovakia) to 0.63 (South Africa) (https://data.worldbank.org/indicator/SI.POV.GINI?end=2024&start=1963&view=chart). For female Standardbreds, Gini indices are 0.694 (Trotters) and 0.733 (Pacers); for males, the values are 0.986 and 0.990, respectively.

### Admixture

Trotters and Pacers have been analyzed separately here, but they don’t represent entirely closed populations. To evaluate the degree of genetic exchange between gaits in our dataset, we computed the fractions of sires and mares that produced foals that were registered under the opposite gait. Consistent with the pattern reported by Cothran et al. (1986), Pacer parents produced same-gait offspring more consistently than did Trotter parents (98.8% for Pacer sires vs 89.4% for Trotter sires, and 98.4% for Pacer dams vs 95.5% for Trotter dams) (Table 1). Foals whose gait differed from a parent subsequently produced fewer offspring than did same-gait foals, and this effect was particularly strong for cross-gait male foals. For foals whose gait differed from the male parent, LRO for male foals on average was only 13-23% of same-gait male foals, and LRO for female foals was 75-94% of same-gait female foals (Table 1). For foals whose gait differed from the female parent, corresponding values were 18-50% for male foals and 66-77% for female foals. After accounting for reduced reproductive success of cross-gait offspring, genetic contributions to admixture from different sources can be estimated as follows. 10.6% of foals sired by a Trotter male had a Pacer gait. Relative LRO for these foals was only 44% (mean of 13% for males and 75% for females), so the effective percentage of cross-gait offspring was 10.6*0.44 = 4.7%. Corresponding values are 0.7% for Pacer sires, 2.9% for Trotter dams, and 1.0% for Pacer dams.

**Table 1.** Summary of cross-gait reproduction in American Standardbreds. Left columns are results for sires; right columns are for dams. T = Trotter; P = Pacer. A: fractions of foals produced by a parent that have the same gait as the parent (TT or PP) or a different gait (TP or PT, with the parental gait first). B: per-capita lifetime reproductive output (LRO) by foals, sorted by sex of the foal and compatibility with parental gait. C: relative LRO (compared to same-gait foals) of foals whose gait differed from their parent.

|  |  |  |  |  |  |  |
| --- | --- | --- | --- | --- | --- | --- |
| **A** | **Sire** |  |  | **Dam** |  |  |
|  | **Foal** | **P** | **T** | **Foal** | **P** | **T** |
|  | **P** | 0.988 | 0.106 | **P** | 0.984 | 0.045 |
|  | **T** | 0.012 | 0.894 | **T** | 0.016 | 0.955 |
| **B** | **LRO by these foals** |  |  | **LRO by these foals** |  |  |
|  | **Sire/Foal** | **Males** | **Females** | **Dam/Foal** | **Males** | **Females** |
|  | **TT** | 2.51 | 2.05 | **TT** | 2.41 | 1.78 |
|  | **TP** | 0.32 | 1.54 | **TP** | 1.20 | 1.37 |
|  | **PT** | 0.46 | 1.67 | **PT** | 0.34 | 0.98 |
|  | **PP** | 2.01 | 1.78 | **PP** | 1.87 | 1.49 |
| **C** | **Relative LRO of offspring** |  |  | **Relative LRO of offspring** |  |  |
|  | **Sire/Foal** | **Males** | **Females** | **Dam/Foal** | **Males** | **Females** |
|  | **TP** | 0.13 | 0.75 | **TP** | 0.50 | 0.77 |
|  | **PT** | 0.23 | 0.94 | **PT** | 0.18 | 0.66 |

The combined Pacer-Trotter pedigree for all foals born 1970-2014 can be visualized as a principal component plot using the R package randPedPCA (Lee et al. 2025). A plot of the first two principal components (Figure 13) shows two mostly distinct lineages with some overlap.

**Figure 13.**
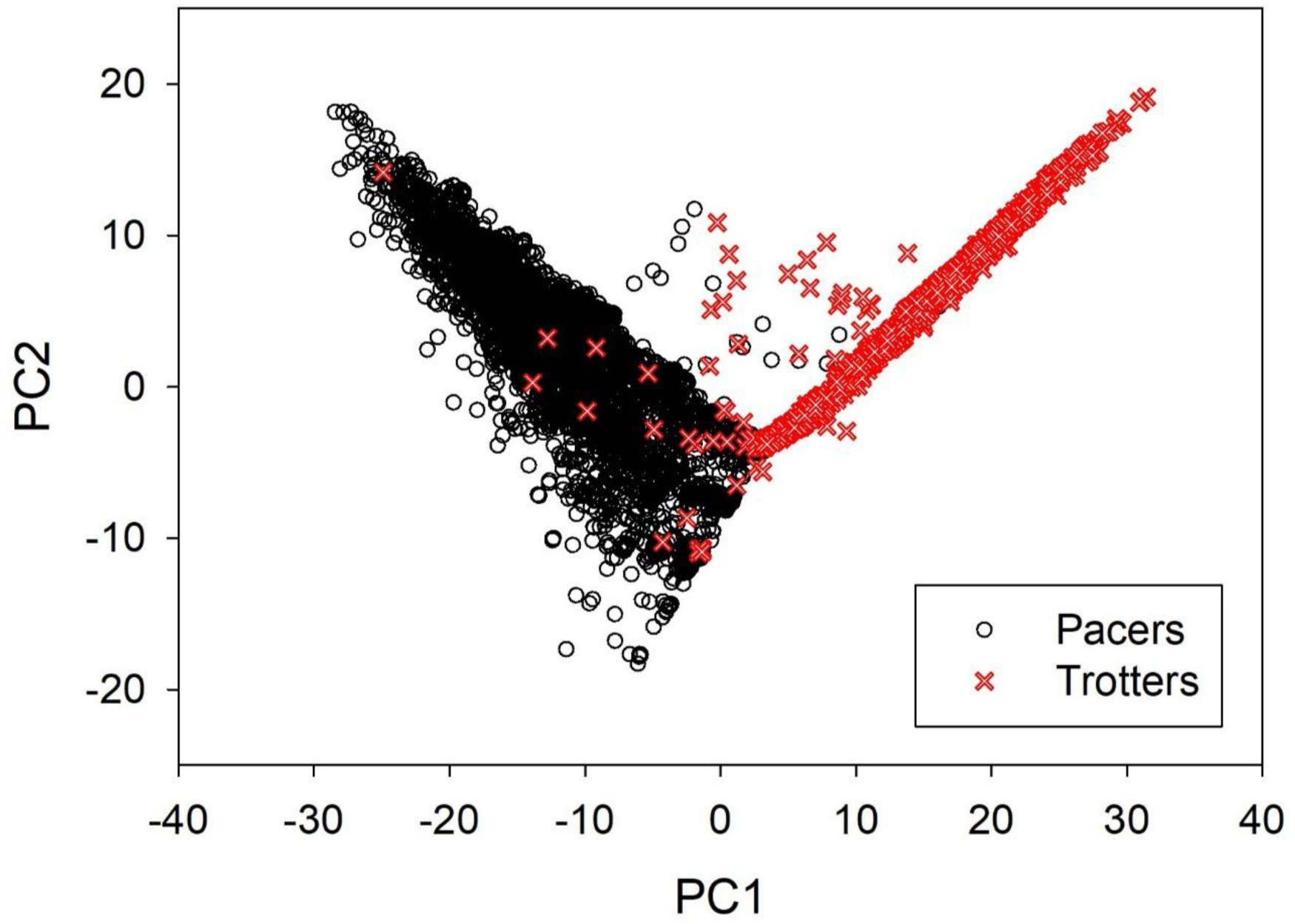
Principal components representation of the combined Pacer-Trotter pedigree for all foals born 1970-2014. Principal components were calculated using the R package randPedPCA described by Lee et a. (2025). PC1 explains 1.2% of the overall variance and PC2 0.7%. The full pedigree contains >700K horses; for this visualization, 10,000 individuals were randomly sampled. In the “mixed” area defined by scores for PC2>2 and –5<PC1<10, most individuals had one Pacer parent and one Trotter parent. PC2 scores were positively correlated (*r* = 0.62; *P* < 0.001) with birth year and hence provide a temporal dimension to the pedigree, with higher PC2 scores associated with more-recently born foals.

## Discussion

Before recapping the major results presented here, we briefly review relevant information that has previously been published for Standardbreds, other horse breeds, and other mammals.

### Previous results for Standardbred horses

Previous pedigree and genetic work on Standardbred horses have focused on inbreeding rather than effective population size. MacCluer et al. (1983) traced pedigrees of up to 30 generations in length. They found that although 1) breeders generally avoid very close inbreeding, and 2) there was no clear evidence for an increase in inbreeding over time, “nearly all horses with complete pedigrees are inbred within the first five ancestral generations” (p. 394). Trotters have higher mean inbreeding coefficients (*F* = 0.103) than Pacers (*F* = 0.074), but “inbreeding does not appear to be a significant factor influencing reproductive performance of Standardbred horses” (Cothran et al. 1984, p. 220; see also Cothran et al. 1986). Higher *F* in Trotters can be explained by the empirical observation (Table 1 and Cothran et al. 1986) that Trotter parents produce foals with Pacer gait more often than the reverse happens. Consequently, Pacers are derived from a more diverse ancestry than Trotters (Cothran et al. 1986).

More recently, Esdaile et al. (2022) used 16 microsatellites to evaluate genetic diversity in Standardbreds, with a focus on the period 2009-2015 (a bit over half a Standardbred generation). They found that genetic differentiation between the gaits (indexed by *F_ST_*) increased (albeit non-significantly) over that period, which could reflect increasingly strong heritability of gait. Departures from Hardy-Weinberg expected genotypic proportions were small (*F_IS_* slightly negative, indicating a small excess of heterozygotes). Esdaile et al. found that, in both gaits, sires who produced more offspring had lower allelic richness, and that sires as a group had lower allelic richness and expected heterozygosity than dams or their offspring. In an exploratory analysis, Esdaile et al. (2022) also reported data for 110K genome-wide SNPs, but these were assayed for only a small subset of sires and their offspring so results are considered provisional.

### Results for other horse breeds

A number of studies have used genetic and/or pedigree methods to estimate *N_e_* in other horse breeds. Genetic studies primarily used LD or coalescent methods, and pedigree studies typically used the rate of increase in inbreeding to estimate effective size. Many of these studies focused on small populations of conservation concern [e.g., Hreiðarsdóttir et al. (2014) Iceland; Takasue et al. (2011) Japan; Duru et al. (2017) Turkey; Maciel et al. (2014) and Madeiros et al. (2024) Brazil; Teegen et al. (2009) Germany; Mousavi et al. (2023) and Bazvand et al. (2024) Iran; Do et al. (2014) Korea; Jasielczuk et al. (2020) Poland; Vázquez-Armijo (2017) Mexico; Stephens et al. (2013) USA]. Not surprisingly, most of the above studies reported low estimates of *N_e_* (at most a few hundred), but few also reported adult census size to allow an estimate of the *N_e_*/*N* ratio. A couple of exceptions were found. In the Do et al. (2014) study of horses on Jejue Island, Korea, the LD-based estimate of recent *N_e_* based on 344K SNPs was 41. This can be compared to a herd size of about 2000, suggesting *N_e_*/*N* of about 0.02. Takasue et al. (2011) estimated contemporary *N_e_* = 48 for the Kiso breed in Japan, compared to a total population size of <150. However, this estimate of *N_e_* only accounted for uneven sex ratio using Wright’s (1938) sex-ratio adjustment, so it undoubtedly is strongly biased upwards. Berg (1986) also used the sex-ratio method to study five isolated populations of feral horses in the U.S. and reported *N_e_*/*N* estimates ranging from 0.2 to 0.6.

Among studies that estimated effective size over time, a Bayesian skyline-plot analysis of the Icelandic horse based on mtDNA (Campana et al. 2011) concluded that *N_e_* had been relatively stable at ∼10K for the last 150 years. In the UK, Corbin et al. (2011) assayed 34K SNPs for 817 UK Thoroughbreds and, using LD-based methods, estimated that *N_e_* had ranged from about 80 to 200 over last 100 generations. Neither of these studies attempted to estimate *N_e_*/*N*.

Finally, two papers conducted meta-analyses of pedigree data and effective size estimates for many domesticated breeds of horses and other livestock species, including cattle, sheep, pigs, goats, and dogs. Leroy et al. (2013) extracted pedigree data from French national databases, and Hall (2016) extracted global information from published and unpublished reports and FAO databases. For 20 French horse breeds, Leroy et al. (2013) found lower estimates using inbreeding methods (mean 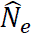 =125-184) than for Hill’s method based on *LRO* (mean 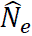 = 487). For 43 global horse breeds, Hall (2016) found smaller estimates for inbreeding-based methods (median 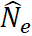 = 66-208) compared to LD methods (median 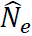 = 194-399). In both studies, effective size estimates for horse breeds were roughly comparable to those for other livestock species. Hall’s review also was able to estimate *N_e_*/*N* for a subset of studies for which census data were available from FAO, and medians for domesticated horses ranged from 0.015 to 0.037, depending on the method.

### Results for other mammals

Two large meta-analyses have tabulated estimates of the *N_e_*/*N* ratio for diverse taxa. Median estimates for mammals were 0.31 as reported by Hoban et al. (2020) and 0.17 as reported by Clarke et al. (2024), with the latter based entirely on estimates using the LD method.

### New results

Results presented here regarding the distribution of reproductive success are novel in several respects. A useful way to quantify annual reproductive skew is to compare the variance in offspring number to the mean—stratified by age and sex. If (for example), all 6-year-old males act like a mini Wright-Fisher population with random reproductive success, the variance in offspring number should approximately equal the mean, leading to 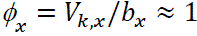. Unfortunately, although many published life tables provide estimates of age-specific fecundity (*b_x_*), few also report the associated variances *V_k,x_* (Waples 2023). The most extreme published estimates of *ϕ_x_* we have seen are for male black bears in Michigan. In this populations, *V_k,x_* rises slowly for young males, being twice the mean at age 3 and three times the mean at age 4, but soon leaps to a high of *ϕ_x_* = 22 at age 7 and a geometric mean of 10.8 for ages 6-10 (Waples et al. 2018b). In Standardbreds, *ϕ* is greater than 10 for all males age 5 or older and exceeds 20 for male Trotters age 25 or older (Table S3). Reproductive skew is even more extreme in male Pacers, with *ϕ* > 40 for all ages 25 and older. Few Standardbred males are still reproductively active after age 20 or so, but most of those that are still active have very high reproductive output.

Like most large mammals, female horses are generally constrained to produce no more than one offspring per year. But a reproductive lifespan of ∼25 years in female Standardbreds allows opportunities for skewed lifetime reproductive success. By 1990, only about 40% of newborn female Trotters would ever produce a registered foal, and for Pacers the fraction was less than 30% (Table S4). Conversely, the most prolific dam within each cohort regularly produced 15-20 foals over its lifetime – compared to a mean of 2 to maintain a stable population. These patterns arise because female reproductive success in Standardbreds is positively correlated over time—females that produce an offspring in a given year are more likely to also produce an offspring the next year (and in subsequent years) than are females that don’t produce an offspring (Figure S2). This pattern can be contrasted with those exhibited by females of many other species who practice skip or intermittent breeding (Goutte et al. 2011; Shaw and Levin 2011, 2013). In these species, reproduction is energetically costly enough that females often skip one or more years after producing an offspring before reproducing again, leading to negative correlations between reproductive output in successive years. In Standardbreds, the positive temporal correlations in female reproductive success lead to values of lifetime variance in offspring number (*V*_*k*•_) that are up to 7 times the mean for Trotters and up to 9 times the mean for Pacers (Table S4; Figure 9).

Male reproductive skew in Standardbreds is much more extreme. Not only do the most prolific males each year sire an astonishingly high number of offspring (Figure 6)—the same males are called upon year after year to produce the next cohort of foals. Positive temporal correlations in foal production in successive years are extremely high (∼0.9) for both Trotter and Pacer males (Figure S2), and these corrwelations generate the most extreme examples of persistent individual differences we have seen in the published literature. As a consequence, lifetime *V*_*k*•_ can exceed 1000 in Trotters and 3000 in Pacers – the latter being 1500 times the mean for a stable population.

Two major factors have facilitated the extreme reproductive skew in male Standardbreds. First, humans rather than the animals themselves decide who mates with whom, and how often. Reproductive dominance by a few males occurs in natural populations of numerous species, especially those with harem polygamy. However, in nature male dominance is generally achieved only through continual, Herculean feats of strength and vigilance. Harem members generally number a few dozen at most, rather than hundreds, and few males can maintain a large harem for long. With Standardbreds, a relatively few males are deemed most desirable by large numbers of horse owners, and these human preferences can persist across lifetimes of favored horses. For Standardbreds, the primary factor that human selection is based upon is racing performance, as well as the ability to produce offspring that are successful at the racetrack.

The second factor is the advent of artificial insemination. In natural horse populations, reproduction is only possible when males and females co-occur in space and time. In a country as large as the U.S., arranging that by human intervention can be costly, logistically challenging, time consuming, and risky. This space/time requirement is obviated by artificial insemination, which relaxes the upper limit to male reproductive output. Artificial insemination has been widely used in Standardbred foal production since the 1950s and in recent years has been responsible for nearly 100% of foal production.

It might seem that gelding, which removes roughly half the males from the gene pool before they reach sexual maturity, is also important for increasing reproductive skew, but we doubt that is actually the case. Of Standardbred males that are NOT gelded, 87% (Trotters) to 88% (Pacers) never sire a foal. It seems likely, therefore, that most males that are gelded are not destined to produce any offspring anyway.

By most measures we investigated, the degree of reproductive skew in Standardbreds increased over time for both sexes in both gaits (Table S5; Figures 5, 7, 9). Starting in 2009 (Trotters) and 2011 (Pacers), the U.S. Trotting Association implemented an annual stud book cap of 140 registered foals per sire (Esdaile et al. 2022). These changes are too recent to have shown up in our data but are not expected to have a large influence on patterns described here, as males that operate within this book cap limit can still sire astronomical numbers of foals across their lifetimes.

High and increasing reproductive variance, especially in males, has predictable consequences for effective population size. Across 21 cohorts, median *N_e_*/*N* ratios were 0.41 and 0.43 for female Trotters and Pacers, respectively (Table S4). This means that although female effectives sizes were still relatively large (medians = 8950 for Trotters and 26687 for Pacers), they were not as large as they could have been if the distribution of *LRO* had been more equitable among females. Male effective sizes were dramatically smaller (medians of 151 for Trotters and 346 for Pacers), and male *N_e_*/*N* was well below 0.01 in both gaits (Table S4). Overall *N_e_* is strongly influenced by the smaller of the sex-specific values. For the most recent 5 cohorts, harmonic means for overall *N_e_* were 570 (Trotters) and 686 (Pacers). These values are just above the minimum threshold of *N_e_*≥500 that Franklin (1980) proposed was necessary to ensure sustainable levels of genetic diversity and well below the minimum size of 1000 suggested by Frankham et al. (2014). These overall *N_e_* values are a small fraction of the total number of adult horses alive at any given time, which were as high as 51K for Trotters and 163K for Pacers (Table S2). For Trotters, the median *N_e_*/*N* ratio across all cohorts was 0.014, and it was below 0.01 for 8 cohorts. In Pacers, *N_e_*/*N* was <0.01 for each of the most recent 8 cohorts and averaged 0.004 for the last 4 cohorts. *N_e_*/*N* ratios smaller than 0.01 are often considered “tiny” and are generally thought to apply primarily to marine species with high fecundity and sweepstakes reproductive success (Hauser and Carvalho 2008; Hedgecock et al. 2011). When *N_e_*/*N* is this low, even large populations can be at risk of losing genetic diversity.

Many of the parameters calculated in this study show a good deal of variation and there is an overall trend of a decline in values over time. The variation can largely be attributed to factors that impact the profitability of raising Standardbreds. Labor cost and worker availability can vary widely, as can costs for feed and medical treatment of horses. Industry-wide effects include competition from alternate forms of gambling and government subsidies within states to support horse breeding efforts. Long term population size declines have resulted from increases in costs associated with horse breeding as well as a decrease in the popularity of horse racing as a form of entertainment across society in the US and other parts of the world.

### Future work

Despite being based on the complete pedigree of U.S. Standardbred horses over 45 years, this study has two limitations. First, it includes no genetic data, which can provide different kinds of insights into inbreeding and its consequences for natural or managed populations. Second, although our study spans about 4.5 horse generations, the moving-window approach we have used with Hill’s model considers only one generation of data at a time (parents and their offspring). Long term effects of inbreeding and reproductive variance depend not only on the distribution of offspring number, but also on the distribution of grandoffspring, great-grandoffspring, etc. Figure 13 does visually represent the multigeneration pedigree, but not directly in terms of reproductive skew and effective population size.

Our future work with these datasets will address these limitations. Microsatellite data are routinely used in Standardbreds to confirm paternity and provide an effective means of identification, although they are rarely used in that manner. The genetic data allow one to track changes in levels of genetic variability, which can be correlated with predictions based on the demographic estimates of *N_e_* reported here. In addition, we will examine evidence for heritability of fitness in Standardbreds. When fitness itself is heritable (that is, when offspring from large families also produce large families), long-term *N_e_* is overestimated by the sequence of single-generation demographic estimates (Robertson 1961; Nei and Murata 1966; Santiago and Caballero 1995). Determining by exactly how much (if at all) multigeneration *N_e_* in Standardbred horses is reduced by this phenomenon is a major objective of our ongoing work.

## Funding and Acknowledgments

Most of the data used in this study was provided by the US Trotting Association and we greatly appreciate their help. USTA personnel also were very helpful in supplying information needed for this study. In particular, Janet Forry, David Carr and T.C. Lane provided tremendous support and this study could not have been completed without them. Portions of this study were funded by a 1yr, $27,325 grant in 2003 from the USTA, “ An analysis of genetic variation in the Standardbred horse.” We thank Tom Reed and Marty Kardos for useful discussions.

## Data Availability

Pedigree data used in this study will be made available on request to the corresponding author.

