## Supporting Info for "Extreme male reproductive skew limits effective population size in American Standardbred horses"

### Supporting Information

Table S1. Yearly numbers of registered Standardbred foals, 1970-2014.

| Year | Pacers | Trotters |
|---|---|---|
| 1970 | 10161 | 4139 |
| 1971 | 10528 | 3983 |
| 1972 | 10693 | 3774 |
| 1973 | 11323 | 3685 |
| 1974 | 11753 | 3967 |
| 1975 | 11990 | 4056 |
| 1976 | 12465 | 4091 |
| 1977 | 13149 | 4379 |
| 1978 | 13955 | 4616 |
| 1979 | 14869 | 4774 |
| 1980 | 16147 | 5294 |
| 1981 | 16702 | 5459 |
| 1982 | 17434 | 5432 |
| 1983 | 17642 | 5777 |
| 1984 | 17864 | 5811 |
| 1985 | 18297 | 6020 |
| 1986 | 17431 | 5956 |
| 1987 | 16970 | 5922 |
| 1988 | 16489 | 5683 |
| 1989 | 15567 | 5318 |
| 1990 | 14765 | 5089 |
| 1991 | 13458 | 4919 |
| 1992 | 12476 | 4545 |
| 1993 | 11809 | 4436 |
| 1994 | 10787 | 4277 |
| 1995 | 10293 | 4160 |
| 1996 | 10070 | 4317 |
| 1997 | 9730 | 4507 |
| 1998 | 9232 | 4730 |
| 1999 | 8996 | 4762 |
| 2000 | 9331 | 5096 |
| 2001 | 9454 | 5278 |
| 2002 | 9562 | 5484 |
| 2003 | 9599 | 5580 |
| 2004 | 9463 | 5775 |
| 2005 | 9299 | 5606 |
| 2006 | 8764 | 5354 |
| 2007 | 8140 | 5020 |
| 2008 | 7472 | 4774 |
| 2009 | 6886 | 4594 |
| 2010 | 6274 | 4352 |
| 2011 | 5908 | 4100 |
| 2012 | 5470 | 4060 |
| 2013 | 4904 | 3635 |
| 2014 | 4121 | 3197 |
| All | 517692 | 215783 |

Table S2. Estimates of total ($N_T$) and adult ($N_A$) abundance by year.

| | Trotters | | Pacers | |
|---|---|---|---|---|
| | $N_T$ | $N_A$ | $N_T$ | $N_A$ |
| 1969 | 55591 | 46673 | 141988 | 119212 |
| 1970 | 54425 | 46673 | 141080 | 119212 |
| 1971 | 53232 | 46673 | 140523 | 119212 |
| 1972 | 51956 | 45626 | 140127 | 118395 |
| 1973 | 50764 | 44554 | 140308 | 117893 |
| 1974 | 49837 | 43408 | 141001 | 117538 |
| 1975 | 49048 | 42336 | 141845 | 117701 |
| 1976 | 48408 | 41501 | 143083 | 118325 |
| 1977 | 48072 | 40793 | 144947 | 119084 |
| 1978 | 48060 | 40224 | 147482 | 120200 |
| 1979 | 48186 | 39931 | 150809 | 121880 |
| 1980 | 48845 | 39928 | 155221 | 124162 |
| 1981 | 49716 | 40052 | 159890 | 127153 |
| 1982 | 50492 | 40654 | 165108 | 131114 |
| 1983 | 51640 | 41487 | 170257 | 135338 |
| 1984 | 52862 | 42270 | 175313 | 140067 |
| 1985 | 54315 | 43385 | 180485 | 144708 |
| 1986 | 55631 | 44559 | 184545 | 149233 |
| 1987 | 57003 | 45935 | 187789 | 153833 |
| 1988 | 58097 | 47180 | 190333 | 157413 |
| 1989 | 58833 | 48458 | 191702 | 160226 |
| 1990 | 59316 | 49467 | 192120 | 162364 |
| 1991 | 59625 | 50118 | 191040 | 163388 |
| 1992 | 59517 | 50521 | 188916 | 163510 |
| 1993 | 59294 | 50738 | 186001 | 162234 |
| 1994 | 58890 | 50549 | 182043 | 159984 |
| 1995 | 58353 | 50269 | 177620 | 157016 |
| 1996 | 57982 | 49828 | 173006 | 153109 |
| 1997 | 57782 | 49247 | 168095 | 148792 |
| 1998 | 57719 | 48807 | 162827 | 144317 |
| 1999 | 57667 | 48526 | 157399 | 139623 |
| 2000 | 57888 | 48383 | 152576 | 134708 |
| 2001 | 58267 | 48258 | 148073 | 129727 |
| 2002 | 58807 | 48402 | 143924 | 125361 |
| 2003 | 59424 | 48723 | 140063 | 121371 |
| 2004 | 60208 | 49200 | 136372 | 117795 |
| 2005 | 60788 | 49767 | 132844 | 114559 |
| 2006 | 61126 | 50511 | 129117 | 111531 |
| 2007 | 61134 | 51080 | 125157 | 108706 |
| 2008 | 60911 | 51439 | 120961 | 105750 |
| 2009 | 60565 | 51501 | 116580 | 102590 |
| 2010 | 59994 | 51334 | 112017 | 99202 |
| 2011 | 59219 | 51038 | 107529 | 95663 |
| 2012 | 58460 | 50536 | 103024 | 91957 |
| 2013 | 57301 | 49833 | 98351 | 88269 |
| 2014 | 55754 | 49121 | 93280 | 84515 |
| Means | 55553 | 46624 | 146426 | 124906 |

Table S3. Age-specific vital rates. $s_x$ = annual survival; $l_x$ = cumulative survival through age $x$; $b_x$ = fecundity; $\phi$ = ratio of variance to mean offspring number.

| | | | Trotters | | | | Pacers | | | |
|---|---|---|---|---|---|---|---|---|---|---|
| | | | Females | | Males | | Females | | Males | |
| Age | $s_x$ | $l_x$ | $b_x$ | $\phi$ | $b_x$ | $\phi$ | $b_x$ | $\phi$ | $b_x$ | $\phi$ |
| **1** | 0.948 | 1.000 | 0 | NA | 0 | NA | 0 | NA | 0 | NA |
| **2** | 0.948 | 0.948 | <0.001 | 1.000 | <0.001 | 7.1 | 0.001 | 0.999 | 0.001 | 13.4 |
| **3** | 0.948 | 0.899 | 0.015 | 0.985 | 0.004 | 7.2 | 0.013 | 0.987 | 0.005 | 12.2 |
| **4** | 0.948 | 0.852 | 0.087 | 0.913 | 0.014 | 9.0 | 0.074 | 0.926 | 0.009 | 11.8 |
| **5** | 0.948 | 0.808 | 0.189 | 0.811 | 0.139 | 10.9 | 0.158 | 0.842 | 0.103 | 13.4 |
| **6** | 0.948 | 0.766 | 0.232 | 0.768 | 0.207 | 12.6 | 0.212 | 0.788 | 0.183 | 13.8 |
| **7** | 0.948 | 0.726 | 0.253 | 0.747 | 0.242 | 12.3 | 0.243 | 0.757 | 0.236 | 12.7 |
| **8** | 0.942 | 0.688 | 0.264 | 0.736 | 0.277 | 12.5 | 0.263 | 0.737 | 0.278 | 13.5 |
| **9** | 0.942 | 0.648 | 0.264 | 0.736 | 0.291 | 13.6 | 0.272 | 0.728 | 0.302 | 14.8 |
| **10** | 0.942 | 0.611 | 0.264 | 0.736 | 0.291 | 12.8 | 0.274 | 0.726 | 0.307 | 14.4 |
| **11** | 0.942 | 0.575 | 0.251 | 0.749 | 0.279 | 12.0 | 0.267 | 0.733 | 0.292 | 14.3 |
| **12** | 0.942 | 0.542 | 0.239 | 0.761 | 0.260 | 12.1 | 0.254 | 0.746 | 0.275 | 14.9 |
| **13** | 0.870 | 0.510 | 0.224 | 0.776 | 0.250 | 12.5 | 0.239 | 0.761 | 0.255 | 17.1 |
| **14** | 0.870 | 0.444 | 0.226 | 0.774 | 0.242 | 11.2 | 0.239 | 0.761 | 0.255 | 16.0 |
| **15** | 0.870 | 0.386 | 0.224 | 0.776 | 0.246 | 14.2 | 0.235 | 0.765 | 0.255 | 12.7 |
| **16** | 0.870 | 0.336 | 0.215 | 0.785 | 0.228 | 15.3 | 0.230 | 0.770 | 0.247 | 15.2 |
| **17** | 0.813 | 0.292 | 0.206 | 0.794 | 0.221 | 13.1 | 0.218 | 0.782 | 0.226 | 15.3 |
| **18** | 0.813 | 0.238 | 0.209 | 0.791 | 0.208 | 16.6 | 0.217 | 0.783 | 0.224 | 18.4 |
| **19** | 0.813 | 0.193 | 0.204 | 0.796 | 0.203 | 16.5 | 0.215 | 0.785 | 0.219 | 23.1 |
| **20** | 0.813 | 0.157 | 0.194 | 0.806 | 0.200 | 16.6 | 0.200 | 0.800 | 0.194 | 22.9 |
| **21** | 0.813 | 0.128 | 0.184 | 0.816 | 0.197 | 18.9 | 0.181 | 0.819 | 0.185 | 27.9 |
| **22** | 0.813 | 0.104 | 0.153 | 0.847 | 0.182 | 16.7 | 0.151 | 0.849 | 0.180 | 29.0 |
| **23** | 0.813 | 0.084 | 0.115 | 0.885 | 0.163 | 16.7 | 0.118 | 0.882 | 0.162 | 28.9 |
| **24** | 0.813 | 0.069 | 0.077 | 0.923 | 0.143 | 15.5 | 0.083 | 0.917 | 0.151 | 38.8 |
| **25** | 0.625 | 0.056 | 0.041 | 0.959 | 0.132 | 17.9 | 0.044 | 0.956 | 0.131 | 46.3 |
| **26** | 0.625 | 0.035 | 0.022 | 0.978 | 0.160 | 22.3 | 0.021 | 0.979 | 0.141 | 44.8 |
| **27** | 0.625 | 0.022 | 0.008 | 0.992 | 0.161 | 21.8 | 0.008 | 0.992 | 0.139 | 55.5 |
| **28** | 0.625 | 0.014 | 0.004 | 0.996 | 0.145 | 27.1 | 0.001 | 0.999 | 0.136 | 43.1 |
| **29** | 0.625 | 0.009 | 0.000 | NA | 0.194 | 24.0 | 0.000 | 1.000 | 0.134 | 51.4 |
| **30** | 0.000 | 0.005 | 0.000 | NA | 0.064 | 24.3 | 0.001 | 0.999 | 0.133 | 62.9 |

Table S4. Metrics of lifetime reproductive success. Results are shown separately for male and female trotters and pacers. Sires/dams: number of members of each cohort that produced at least 1 offspring during its lifetime; %Sires/%dams: fraction of cohort members that ever reproduced; Noff = total foal production by the cohort; Max = maximum lifetime production by one individual; kbar = mean lifetime offspring production; Vk1: raw variance in lifetime reproductive success; G2: generation length for the cohort; Oldest: oldest age at reproduction; Vk2: adjusted variance in lifetime reproductive success; Ne = effective size per generation; Ne divided by estimated $N_A$ from Table S2 at birth of the cohort.

**Trotters**

| Cohort | Males | Sires | Noff | Max | kbar | Vk1 | G2 | Oldest | Vk2 | Ne | Ne/N | %Sires |
|---|---|---|---|---|---|---|---|---|---|---|---|---|
| 1970 | 1646 | 186 | 3077 | 349 | 1.87 | 264 | 10.79 | 26 | 302 | 234 | 0.01 | 0.113 |
| 1971 | 1613 | 196 | 4373 | 1117 | 2.71 | 1050 | 11.52 | 24 | 572 | 130 | 0.0056 | 0.122 |
| 1972 | 1541 | 170 | 3175 | 865 | 2.06 | 614 | 10.29 | 25 | 578 | 109 | 0.0048 | 0.110 |
| 1973 | 1567 | 187 | 3362 | 734 | 2.15 | 489 | 11.26 | 28 | 425 | 165 | 0.0074 | 0.119 |
| 1974 | 1600 | 199 | 7856 | 1434 | 4.91 | 2249 | 11.68 | 30 | 374 | 199 | 0.0092 | 0.124 |
| 1975 | 1676 | 176 | 6085 | 880 | 3.63 | 1391 | 10.66 | 26 | 423 | 168 | 0.0079 | 0.105 |
| 1976 | 1700 | 149 | 2282 | 631 | 1.34 | 299 | 9.77 | 27 | 663 | 100 | 0.0048 | 0.088 |
| 1977 | 1921 | 165 | 3276 | 1083 | 1.71 | 688 | 10.68 | 26 | 945 | 87 | 0.0042 | 0.086 |
| 1978 | 2030 | 171 | 5635 | 1791 | 2.78 | 2077 | 12.16 | 26 | 1079 | 91 | 0.0045 | 0.084 |
| 1979 | 2093 | 160 | 7749 | 794 | 3.70 | 1469 | 9.61 | 27 | 430 | 186 | 0.0093 | 0.076 |
| 1980 | 2330 | 196 | 6272 | 585 | 2.69 | 483 | 9.74 | 28 | 267 | 337 | 0.0169 | 0.084 |
| 1981 | 2439 | 158 | 6599 | 932 | 2.71 | 1196 | 10.85 | 28 | 654 | 161 | 0.0081 | 0.065 |
| 1982 | 2530 | 172 | 5000 | 599 | 1.98 | 455 | 10.70 | 25 | 466 | 231 | 0.0114 | 0.068 |
| 1983 | 2603 | 146 | 4051 | 1013 | 1.56 | 611 | 10.90 | 25 | 1009 | 112 | 0.0054 | 0.056 |
| 1984 | 2686 | 119 | 3073 | 691 | 1.14 | 287 | 12.14 | 25 | 875 | 149 | 0.007 | 0.044 |
| 1985 | 2844 | 117 | 3099 | 535 | 1.09 | 291 | 9.71 | 24 | 980 | 112 | 0.0052 | 0.041 |
| 1986 | 2810 | 129 | 3327 | 471 | 1.18 | 207 | 11.31 | 28 | 590 | 215 | 0.0096 | 0.046 |
| 1987 | 2863 | 100 | 3576 | 453 | 1.25 | 254 | 10.77 | 24 | 649 | 189 | 0.0082 | 0.035 |
| 1988 | 2740 | 85 | 4067 | 797 | 1.48 | 540 | 10.06 | 23 | 979 | 112 | 0.0048 | 0.031 |
| 1989 | 2632 | 115 | 6187 | 1119 | 2.35 | 1041 | 10.84 | 25 | 754 | 151 | 0.0062 | 0.044 |
| 1990 | 2455 | 95 | 6007 | 1235 | 2.45 | 1491 | 11.31 | 24 | 996 | 111 | 0.0045 | 0.039 |

| Cohort | Females | Dams | Noff | Max | kbar | Vk1 | G2 | Oldest | Vk2 | Ne | Ne/N | % Dams |
|---|---|---|---|---|---|---|---|---|---|---|---|---|
| 1970 | 1766 | 1265 | 4245 | 21 | 2.40 | 8.85 | 11.37 | 26 | 6.46 | 9492 | 0.4068 | 0.716 |
| 1971 | 1711 | 1274 | 4572 | 18 | 2.67 | 9.97 | 11.24 | 26 | 6.09 | 9516 | 0.4078 | 0.745 |
| 1972 | 1638 | 1158 | 4262 | 16 | 2.60 | 10.31 | 11.05 | 26 | 6.56 | 8462 | 0.3709 | 0.707 |
| 1973 | 1629 | 1215 | 4503 | 17 | 2.76 | 10.02 | 10.77 | 27 | 5.80 | 9000 | 0.404 | 0.746 |
| 1974 | 1799 | 1258 | 4880 | 18 | 2.71 | 10.87 | 10.70 | 27 | 6.43 | 9130 | 0.4206 | 0.699 |
| 1975 | 1814 | 1305 | 5088 | 17 | 2.80 | 10.92 | 10.47 | 26 | 6.13 | 9348 | 0.4416 | 0.719 |
| 1976 | 1898 | 1242 | 4891 | 17 | 2.58 | 9.99 | 10.19 | 26 | 6.47 | 9136 | 0.4403 | 0.654 |
| 1977 | 1947 | 1210 | 4662 | 18 | 2.39 | 10.40 | 10.20 | 26 | 7.58 | 8290 | 0.4065 | 0.622 |
| 1978 | 2139 | 1313 | 5059 | 18 | 2.37 | 9.97 | 9.96 | 27 | 7.44 | 9031 | 0.449 | 0.614 |
| 1979 | 2210 | 1306 | 5118 | 17 | 2.32 | 10.19 | 10.02 | 28 | 7.87 | 8979 | 0.4497 | 0.591 |

| | | | | | | | | | | | | |
|---|---|---|---|---|---|---|---|---|---|---|---|---|
| 1980 | 2508 | 1359 | 5446 | 18 | 2.17 | 10.64 | 10.21 | 25 | 9.19 | 9154 | 0.4585 | 0.542 |
| 1981 | 2639 | 1316 | 5324 | 19 | 2.02 | 9.79 | 10.03 | 26 | 9.64 | 9091 | 0.454 | 0.499 |
| 1982 | 2494 | 1191 | 4841 | 16 | 1.94 | 9.48 | 10.04 | 27 | 10.00 | 8344 | 0.4105 | 0.478 |
| 1983 | 2787 | 1184 | 5330 | 20 | 1.91 | 10.70 | 10.36 | 25 | 11.61 | 8489 | 0.4093 | 0.425 |
| 1984 | 2796 | 1185 | 5002 | 17 | 1.79 | 9.64 | 10.49 | 27 | 11.81 | 8494 | 0.4019 | 0.424 |
| 1985 | 2889 | 1054 | 4710 | 18 | 1.63 | 9.31 | 10.40 | 26 | 13.56 | 7721 | 0.3559 | 0.365 |
| 1986 | 2827 | 896 | 4138 | 16 | 1.46 | 8.28 | 10.56 | 25 | 14.73 | 7135 | 0.3203 | 0.317 |
| 1987 | 2861 | 947 | 4243 | 16 | 1.48 | 8.15 | 10.69 | 26 | 14.12 | 7590 | 0.3305 | 0.331 |
| 1988 | 2751 | 960 | 4556 | 17 | 1.66 | 9.29 | 10.64 | 25 | 13.13 | 7735 | 0.3279 | 0.349 |
| 1989 | 2538 | 948 | 4494 | 19 | 1.77 | 9.44 | 10.68 | 25 | 11.78 | 7869 | 0.3248 | 0.374 |
| 1990 | 2493 | 1039 | 5201 | 15 | 2.09 | 10.56 | 10.58 | 24 | 9.79 | 8950 | 0.3618 | 0.417 |

### Pacers

| Cohort | Males | Sires | Noff | Max | kbar | Vk1 | G2 | Oldest | Vk2 | Ne | Ne/N | %Sires |
|---|---|---|---|---|---|---|---|---|---|---|---|---|
| 1970 | 5561 | 498 | 10782 | 721 | 1.94 | 396 | 11.46 | 27 | 422 | 602 | 0.0101 | 0.085 |
| 1971 | 5661 | 509 | 11372 | 883 | 2.01 | 442 | 11.23 | 26 | 438 | 578 | 0.0097 | 0.084 |
| 1972 | 5706 | 479 | 15003 | 1712 | 2.63 | 967 | 10.69 | 26 | 560 | 434 | 0.0073 | 0.081 |
| 1973 | 5933 | 489 | 16679 | 1249 | 2.81 | 1215 | 10.17 | 26 | 615 | 391 | 0.0066 | 0.076 |
| 1974 | 6181 | 488 | 22806 | 1989 | 3.69 | 2161 | 10.61 | 29 | 636 | 411 | 0.007 | 0.071 |
| 1975 | 6334 | 504 | 16613 | 1805 | 2.62 | 1374 | 11.22 | 26 | 799 | 355 | 0.006 | 0.072 |
| 1976 | 6489 | 470 | 17080 | 1982 | 2.63 | 1330 | 10.27 | 30 | 768 | 346 | 0.0058 | 0.069 |
| 1977 | 6868 | 439 | 17948 | 1752 | 2.61 | 1451 | 10.22 | 29 | 850 | 329 | 0.0055 | 0.059 |
| 1978 | 7246 | 408 | 13121 | 912 | 1.81 | 584 | 9.60 | 28 | 712 | 390 | 0.0065 | 0.050 |
| 1979 | 7612 | 442 | 18222 | 2541 | 2.39 | 1527 | 10.42 | 28 | 1066 | 297 | 0.0049 | 0.054 |
| 1980 | 8179 | 428 | 13867 | 796 | 1.70 | 389 | 9.34 | 26 | 541 | 563 | 0.0091 | 0.047 |
| 1981 | 8430 | 409 | 13904 | 1338 | 1.65 | 644 | 10.26 | 28 | 946 | 365 | 0.0057 | 0.044 |
| 1982 | 9026 | 412 | 15063 | 1063 | 1.67 | 670 | 9.88 | 26 | 961 | 370 | 0.0057 | 0.041 |
| 1983 | 8979 | 331 | 14769 | 1807 | 1.64 | 1075 | 11.48 | 30 | 1589 | 259 | 0.0038 | 0.034 |
| 1984 | 9167 | 312 | 11149 | 1446 | 1.22 | 762 | 10.58 | 25 | 2059 | 188 | 0.0027 | 0.032 |
| 1985 | 9274 | 304 | 11705 | 1329 | 1.26 | 672 | 11.40 | 26 | 1686 | 250 | 0.0035 | 0.030 |
| 1986 | 8858 | 257 | 9482 | 1013 | 1.07 | 333 | 10.03 | 24 | 1162 | 305 | 0.0041 | 0.026 |
| 1987 | 8571 | 234 | 9702 | 2321 | 1.13 | 917 | 11.94 | 27 | 2861 | 143 | 0.0019 | 0.024 |
| 1988 | 8291 | 214 | 10603 | 1710 | 1.28 | 968 | 10.77 | 26 | 2366 | 151 | 0.0019 | 0.024 |
| 1989 | 7933 | 217 | 12220 | 1965 | 1.54 | 1427 | 12.44 | 25 | 2405 | 164 | 0.002 | 0.025 |
| 1990 | 7361 | 225 | 8519 | 1423 | 1.16 | 619 | 10.68 | 24 | 1848 | 170 | 0.0021 | 0.027 |

| Cohort | Females | Dams | Noff | Max | kbar | Vk1 | G2 | Oldest | Vk2 | Ne | Ne/N | % Dams |
|---|---|---|---|---|---|---|---|---|---|---|---|---|
| 1970 | 5223 | 3410 | 14803 | 19 | 2.83 | 11.51 | 11.14 | 27 | 6.32 | 27963 | 0.4691 | 0.658 |
| 1971 | 5427 | 3487 | 15294 | 19 | 2.82 | 11.59 | 10.98 | 26 | 6.42 | 28302 | 0.4748 | 0.649 |
| 1972 | 5515 | 3389 | 14724 | 19 | 2.67 | 10.98 | 10.68 | 26 | 6.67 | 27178 | 0.4591 | 0.616 |
| 1973 | 5844 | 3564 | 15615 | 20 | 2.67 | 11.32 | 10.61 | 27 | 6.85 | 28040 | 0.4757 | 0.609 |
| 1974 | 6117 | 3686 | 16052 | 19 | 2.62 | 10.87 | 10.31 | 26 | 6.79 | 28693 | 0.4882 | 0.599 |
| 1975 | 6151 | 3604 | 15452 | 17 | 2.51 | 10.58 | 10.05 | 27 | 7.12 | 27124 | 0.4609 | 0.582 |
| 1976 | 6433 | 3694 | 15749 | 19 | 2.45 | 10.53 | 9.99 | 26 | 7.39 | 27368 | 0.4626 | 0.572 |
| 1977 | 6745 | 3756 | 15546 | 18 | 2.30 | 9.98 | 9.73 | 25 | 7.78 | 26830 | 0.4506 | 0.554 |

| | | | | | | | | | | | | |
|---|---|---|---|---|---|---|---|---|---|---|---|---|
| 1978 | 7131 | 3713 | 15037 | 18 | 2.11 | 9.10 | 9.63 | 26 | 8.29 | 26687 | 0.444 | 0.517 |
| 1979 | 7688 | 3900 | 15977 | 18 | 2.08 | 9.52 | 9.75 | 26 | 8.89 | 27546 | 0.452 | 0.500 |
| 1980 | 8376 | 3895 | 15730 | 19 | 1.88 | 8.97 | 9.67 | 26 | 10.04 | 26910 | 0.4335 | 0.465 |
| 1981 | 8613 | 3680 | 14732 | 18 | 1.71 | 8.30 | 9.70 | 26 | 11.01 | 25689 | 0.4041 | 0.423 |
| 1982 | 8811 | 3505 | 13877 | 20 | 1.58 | 7.90 | 9.77 | 26 | 12.19 | 24274 | 0.3703 | 0.393 |
| 1983 | 9031 | 3282 | 13303 | 20 | 1.47 | 7.81 | 9.87 | 26 | 13.68 | 22737 | 0.336 | 0.359 |
| 1984 | 9006 | 2947 | 11969 | 21 | 1.33 | 7.21 | 9.89 | 26 | 15.32 | 20574 | 0.2938 | 0.323 |
| 1985 | 9275 | 2866 | 11772 | 17 | 1.27 | 6.93 | 10.07 | 26 | 16.06 | 20693 | 0.286 | 0.305 |
| 1986 | 8870 | 2522 | 10755 | 19 | 1.21 | 7.04 | 10.24 | 25 | 17.87 | 18279 | 0.245 | 0.285 |
| 1987 | 8579 | 2266 | 9555 | 18 | 1.11 | 6.35 | 10.19 | 26 | 18.88 | 16743 | 0.2177 | 0.262 |
| 1988 | 8371 | 2231 | 9769 | 18 | 1.17 | 6.67 | 10.42 | 26 | 18.17 | 17307 | 0.2199 | 0.264 |
| 1989 | 7747 | 2086 | 9138 | 18 | 1.18 | 6.74 | 10.41 | 25 | 17.98 | 16147 | 0.2016 | 0.265 |
| 1990 | 7530 | 2032 | 9264 | 16 | 1.23 | 7.10 | 10.38 | 24 | 17.51 | 16017 | 0.1973 | 0.267 |

Table S5. Results of statistical tests for linear trends. Tests for annual trends used data for 45 years; tests for lifetime trends used data for 21 cohorts. Signs (+ or -) indicate the direction for significant trends. * $p<0.05$; ** $p<0.01$; *** $p<0.001$. For annual data, trends were evaluated for numbers of foals and sires; maximum foal production by one stud; effective number of male breeders and the $N_b/N$ ratio for males; the percent of males that sired at least one foal; and the percentage of all foals sired by the most prolific male. For lifetime data, trends were evaluated separately by sex for total number of foals produced by each cohort ($N_1$); maximum foal production by one individual; lifetime variance in offspring number; effective size per generation and the Ne/N ratio; and the percent ... .

**Annual**

| | Trotters | Pacers |
|---|---|---|
| Foals | ns | (-)*** |
| Sires | (-)*** | (-)*** |
| Max | (+)** | ns |
| Male$N_b$ | (-)*** | (-)*** |
| $N_b/N$ | (-)*** | (-)*** |
| PctSires | (-)*** | (-)*** |
| MaxPct | (+)** | (+)*** |

**Lifetime**

| | Trotters | | Pacers | |
|---|---|---|---|---|
| | Males | Females | Males | Females |
| $N_1$ | (+)*** | (+)*** | (+)*** | (+)*** |
| Max | ns | ns | ns | ns |
| $V_k$* | (+)* | (+)*** | (+)*** | (+)*** |
| $N_e$ | ns | (-)*** | (-)*** | (-)*** |
| $N_e/N$ | ns | (-)** | (-)*** | (-)*** |
| Pct | (-)*** | (-)*** | (-)*** | (-)*** |

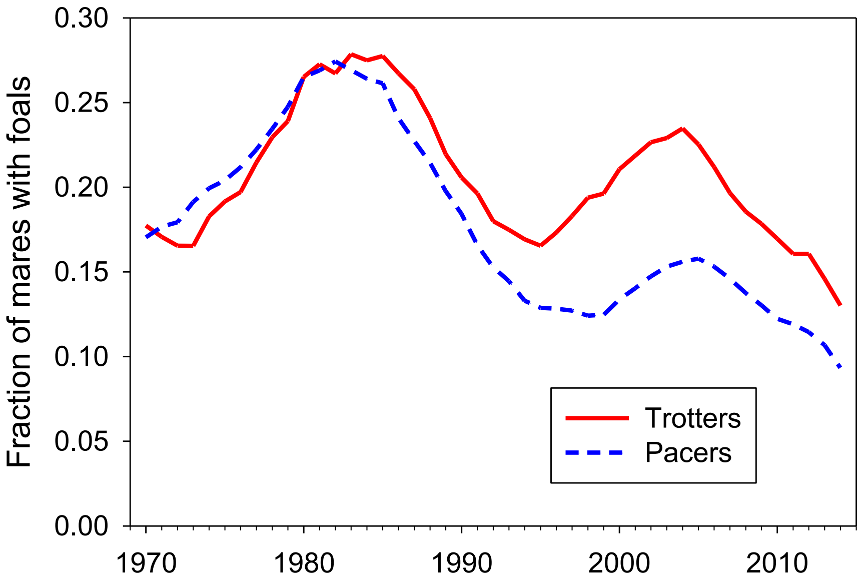

Figure S1. Estimated fraction of adult females that produced a foal each year, 1970-2014.

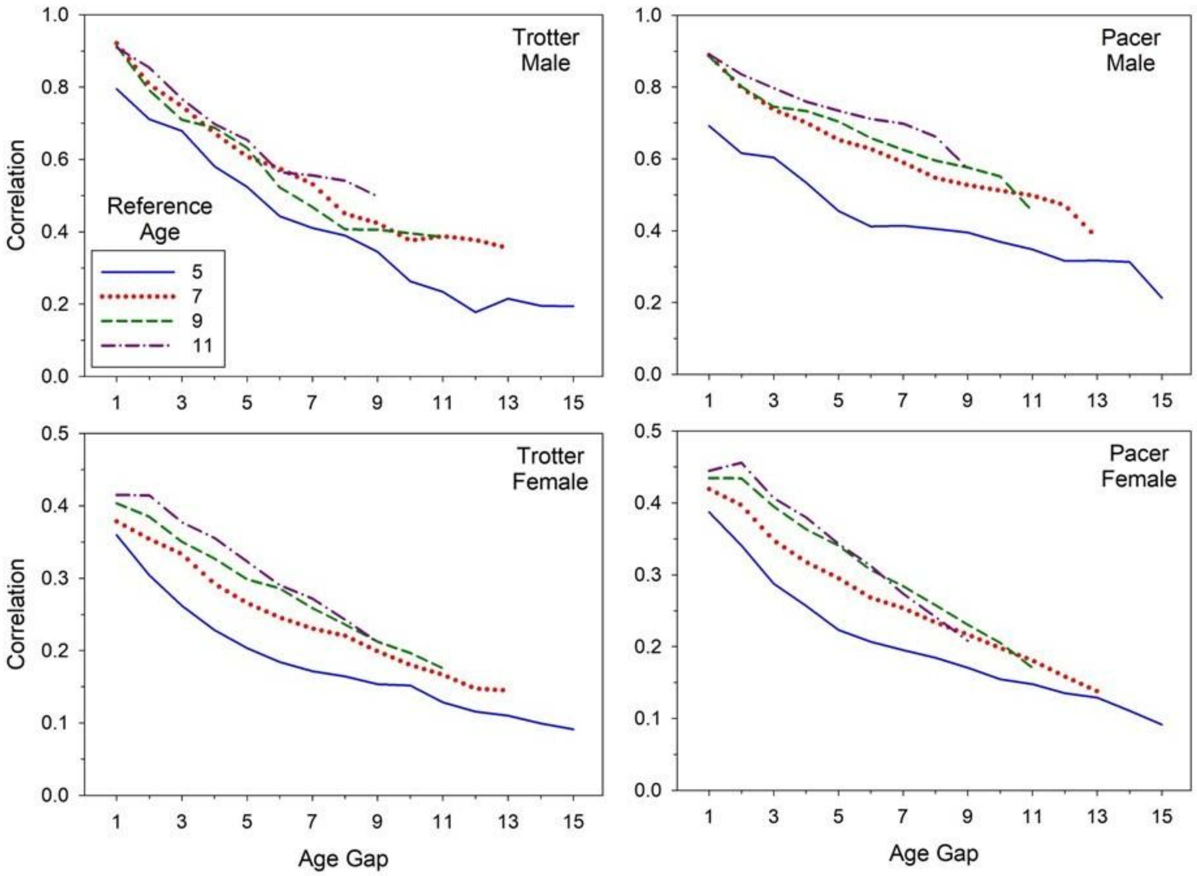

Figure S2. Correlations between number of offspring produced by individuals at different ages. Top panels show results for males, and bottom panels show results for females. Left panels are for Trotters and right panels for Pacers. The X axis shows the gap (in years) between the two ages being compared. Within each panel, results are shown for 4 different reference ages (5,7,9,11), which is the younger of two ages being compared. For example, for reference age = 5 (solid blue lines), results are shown for comparison of reproductive output at ages 5 and 6 (for age gap = 1) through ages 5 and 20 (age gap = 15). Note the different Y axis scales for males and females.
